# Microglial P2Y_6_ calcium signaling promotes phagocytosis and shapes neuroimmune responses in epileptogenesis

**DOI:** 10.1101/2023.06.12.544691

**Authors:** Anthony D. Umpierre, Bohan Li, Katayoun Ayasoufi, Shunyi Zhao, Manling Xie, Grace Thyen, Benjamin Hur, Jiaying Zheng, Yue Liang, Zhaofa Wu, Xinzhu Yu, Jaeyun Sung, Aaron J. Johnson, Yulong Li, Long-Jun Wu

**Author notes:** Correspondence (A.D.U.), (Y.L.), (L-J. W.).

## Abstract

Microglial calcium signaling is rare in a baseline state but shows strong engagement during early epilepsy development. The mechanism and purpose behind microglial calcium signaling is not known. By developing an *in vivo* UDP fluorescent sensor, GRAB_UDP1.0_, we discovered that UDP release is a conserved response to seizures and excitotoxicity across brain regions. UDP signals to the microglial P2Y_6_ receptor for broad increases in calcium signaling during epileptogenesis. UDP-P2Y_6_ signaling is necessary for lysosome upregulation across limbic brain regions and enhances production of pro-inflammatory cytokines—TNFα and IL-1β. Failures in lysosome upregulation, observed in P2Y_6_ KO mice, can also be phenocopied by attenuating microglial calcium signaling in Calcium Extruder (“CalEx”) mice. In the hippocampus, only microglia with P2Y_6_ expression can perform full neuronal engulfment, which substantially reduces CA3 neuron survival and impairs cognition. Our results demonstrate that calcium activity, driven by UDP-P2Y_6_ signaling, is a signature of phagocytic and pro-inflammatory function in microglia during epileptogenesis.

## INTRODUCTION

As innate immune cells in the central nervous system (CNS), microglia play a critical early role in coordinating neuroimmune responses^1,2^. Microglial patterns of calcium activity are fundamentally different from local astrocytes and neurons. In a naïve (non-pathological) baseline state, microglial rarely display spontaneous calcium transients^3-5^. However, microglia begin to transduce calcium signaling following CNS injury, inflammation, or hyperactivity^4-7^. Therein, microglial calcium signaling may reflect the engagement of pathway(s) selectively utilized during shifts away from homeostasis.

The early phase of epilepsy development, or epileptogenesis, represents an ideal context to study the mechanism(s) driving microglial calcium signaling and their role in injury responses and disease progression. Epileptogenesis, when modelled by the systemic administration of kainate, induces a prolonged seizure state known as status epilepticus (SE or KA-SE), which results in cell death^8-10^, reactive gliosis^8,9^, and network rearrangement^11,12^. In the days following SE, microglia display prolonged, whole-cell calcium transients^5^, which are markedly distinct from calcium signaling displayed in the quiescent state.

We hypothesized that microglial calcium elevations in epileptogenesis are driven by purinergic signaling. Purines are released during periods of damage, inflammation, and hyperexcitability, with the ability to evoke calcium signaling in most cell types^13-18^. In glia, large calcium transients are often ascribed to metabotropic G_αq_-PLC-IP_3_ pathway activation, such as the purinergic P2Y_1_ receptor^17^, metabotropic glutamate receptor 5 (mGluR_5_)^19-21^, and α1-noradrenergic receptor (α1AR)^22^ systems in astrocytes. According to transcriptomic databases^23^, only one purine-sensitive receptor, P2Y_6_, is linked to the intracellular G_αq_-PLC-IP_3_ calcium pathway in microglia. The P2Y_6_ receptor is highly enriched in microglia^24^. In the brain, its expression is restricted to microglia and a subset of hypothalamic neurons^25,26^. P2Y_6_ has its highest endogenous affinity for the purine UDP, which is known to be elevated during epileptogenesis^25^.

We investigated how UDP-P2Y_6_ signaling influences microglial calcium signaling and function during epileptogenesis. To visualize real-time UDP signaling in the mouse brain, we developed GRAB_UDP1.0_ (abbreviated UDP1.0), a novel UDP fluorescent sensor. Using two-photon imaging, we find evidence for enhanced UDP release in hippocampus and cortex during early epileptogenesis, which coincides with increased UDP calcium sensitivity in microglia. Abolishing microglial UDP responses through P2Y_6_ knockout results in a strong, longitudinal attenuation of spontaneous microglial calcium signaling during epileptogenesis. Furthermore, transcriptomics, histology, and high-parameter flow cytometry suggest that the salient emergence of microglial calcium signaling in early epileptogenesis coordinates pro-inflammatory and pro-phagocytic state changes, which have an adverse effect on pyramidal neuron survival and hippocampal cognition.

## RESULTS

### UDP evokes calcium activity in microglia, with enhanced sensitivity during epileptogenesis

UDP is a strong candidate molecule for activating larger-scale calcium signaling in microglia, based upon its engagement of the G_αq_-linked P2Y_6_ receptor. Indeed, even in naïve microglia, we observe robust calcium responses to UDP in the range of 100 µM to 1 mM (Cx3Cr1^CreER-IRES-eYFP^; R26^LSL-CAG-GCaMP6s^ acute brain slice; Figures S1A-S1C). In this same concentration range, ATP calcium responses are markedly smaller in magnitude (Figures S1A-S1C). UDP calcium responses have a prolonged duration (vs. ATP calcium responses; Figure S1D), which may have relevance for the multi-minute calcium events we have previously reported *in vivo*^5^. UDP application predominately transduces calcium signaling in microglia via P2Y_6_ receptors, as the selective antagonist MRS-2578 (40 µM) reduces UDP responses by 82.1 ± 5.8% (vs. 3.2 ± 10.9% attenuation due to desensitization; Figures S1E-S1G). UDP has minimal calcium activity at other purine receptors, including P2X_4_/P2X_7_, and P2Y_12_, based on antagonist studies (Figures S1F and S1G). Pharmacological studies therefore suggest that UDP could work at micromolar levels to promote calcium activity in microglia via P2Y_6_ signaling.

P2Y_6_ expression in microglia is often up-regulated following brain insults^24^, which may contribute to enhanced microglial calcium signaling during epileptogenesis. We assessed heightened P2Y_6_ receptor expression based upon the magnitude of UDP calcium responses (Figure S2A). This sensitivity-based approach was chosen because of its direct interpretability and difficulties with P2Y_6_ commercial antibody interpretability^27^. At baseline, approximately 50% of microglia respond to UDP in a naïve state (Figures S2B, S2D, and S2F), matching the observation of P2Y_6_ enrichment in microglia^24,25^. Following KA-SE, there are brain-wide enhancements in microglial UDP calcium responses. In cortex, UDP sensitivity is elevated across all epileptogenesis time points (Day 1, 3, 7, and 10), with a maximal 40-60% increase in calcium responses during the first week (Figure S2C). In the hippocampus, UDP sensitivity is also elevated across all time points, but shows a pattern of ever-increasing sensitivity over time. In CA3 and CA1, UDP calcium responses double 10 days after KA-SE (96% increase in CA3, 118% increase in CA1; Figures S2E and S2G). In addition, a greater proportion of microglia begin to respond to UDP in the hippocampus during epileptogenesis (from approximately 50% at baseline to 75% from Day 3-10; Figures S2D and S2F). These results suggest that the P2Y_6_ receptor has heightened sensitivity to UDP during epileptogenesis, evidenced through its enhanced calcium mobilization in microglia.

### Development of a fluorescent sensor for *in vivo* imaging of extracellular UDP

*Ex vivo* studies demonstrate that UDP can evoke microglial calcium signaling. On the other hand, whether UDP serves as an endogenous activator of microglial calcium signaling during epileptogenesis requires *in vivo* knowledge of its temporal and spatial release patterns. To understand endogenous UDP release dynamics, we developed and characterized a high-performance, GPCR-activation-based (GRAB) sensor for UDP. To achieve this, we inserted a circularly permutated, enhanced GFP (cpEGFP) molecule, with flanking linker sequences derived from GRAB_NE1m_^28^, between transmembrane domains 5 and 6 of the chicken P2Y_6_ (cP2Y_6_) receptor, forming a modified intracellular loop (Figure 1A). By optimizing the length and residue composition of the linker sequences, we produced a candidate sensor with strong fluorescent responses to UDP, which we termed GRAB_UDP1.0_, or UDP1.0 (Figures 1B, S3A, and S3B). UDP1.0 is properly trafficked to the plasma membrane (Figure S3C) and displays an approximate 6-fold ΔF/F response to 10 µM UDP (Figures 1C and 1D). Its single-photon excitation and emission spectra are similar to EGFP (Figure 1E). Non-selective P2Y receptor antagonists, Suramin and PPADS, can disrupt UDP1.0 fluorescent responses to UDP (Figure 1F). The sensor also displays strong selectivity for UDP over other purines and neurotransmitters (Figure 1F). After UDP, UTP has the next highest affinity for the sensor with a 16-fold difference in EC_50_ value (91.4 nM UDP vs. 1.28 µM UTP; Figure 1G). We additionally performed a split-luciferase complementation assay to examine whether UDP1.0 would couple to a native G_αq_ pathway (Figure 1H). UDP1.0 exhibits minimal G_αq_-dependent coupling, similar to non-transfected control cells, while the wild-type cP2Y_6_ receptor can induce robust Gq-coupling as reported by downstream luciferase luminescence (Figure 1H). Likewise, a red-shifted calcium indicator dye reports calcium mobilization in HEK293T cells transfected with the cP2Y_6_ receptor, but not with the UDP1.0 construct (Figure S3D-S3F). These results suggest that UDP1.0 expression does not significantly couple with native G_αq_ proteins and is therefore unlikely to induce robust G_αq_ signaling in cell types that were not otherwise responsive to UDP. In addition, the sensor is unlikely to undergo internalization or experience other mechanisms reducing sensitivity over time as 2-hour incubation with 100 µM UDP does not significantly reduce sensor fluorescence over time (Figure 1I). Therefore, *in vitro* characterization demonstrates that UDP1.0 can detect UDP in the nanomolar to micromolar range with strong selectivity for this purine over all other ligands tested.

**Figure 1:**
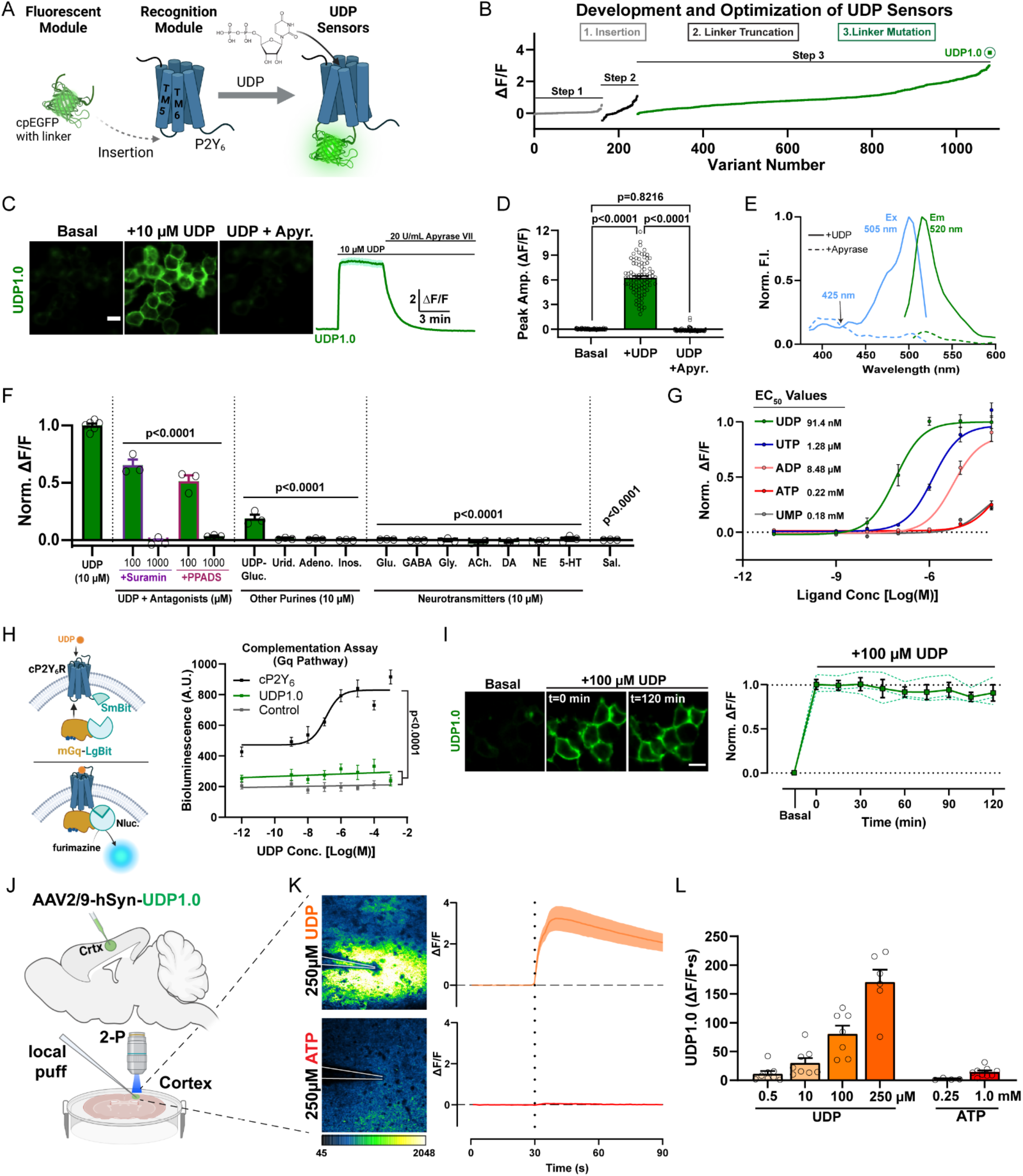
Development of a fluorescent sensor for *in vivo* imaging of extracellular UDP. (A) To engineer the UDP biosensor, a circularly permutated enhanced GFP (cpEGFP) molecule with linker sequences was inserted into the extracellular loop between transmembrane (TM) 5 and 6 of the chicken P2Y_6_ (cP2Y_6_) receptor. (B) Optimization of the UDP sensor occurred through a three-step process including insertion site variation, N- and C-terminal linker truncation, and point-mutation-based optimization of the linker sequence, yielding the UDP1.0 variant (see Figures S3A, S3B). (C) Representative images and corresponding ΔF/F trace from HEK293T cells expressing UDP1.0 under basal conditions, after the addition of 10 µM UDP, and then after 20 U/mL apyrase grade VII (apyr.) treatment. Scale bar, 20 µm. (D) Summary of peak UDP1.0 ΔF/F amplitude under basal, UDP, or UDP plus apyrase conditions. One-Way ANOVA with Tukey’s post-hoc test (n=90 total cellular ROIs). (E) One-photon excitation (Ex) and emission (Em) spectra for UDP1.0 in the presence of 10 µM UDP (solid lines), or 5 U/mL apyrase (dashed lines). (F and G) Selectivity of UDP1.0. (F) Comparison of UDP responses in the presence of non-selective P2Y receptor antagonists, or other purine agonists, neurotransmitters, and saline. Peak ΔF/F responses were normalized to UDP. One-Way ANOVA with post-hoc UDP comparison. (G) UDP1.0 fluorescent response curves to purinergic ligands with EC_50_ values, see Star Methods for fitting (n=3-6 wells; 300-500 cells per well). (H) Wild-type cP2Y_6_, but not UDP1.0, drives Gαq signaling measured using a luciferase complementation assay. Left, schematic of the complementation assay. Right, total luminescence emitted by cells co-transfected with LgBit-mGq alone (control), or LgBit-mGq with cP2Y_6_-SmBit, or UDP1.0-SmBit, plotted against UDP concentration. One-Way ANOVA with Tukey’s post-hoc test (UDP1.0 vs. Control: p=0.3406; n=3 wells/group). (I) UDP1.0 fluorescence in HEK cells under basal conditions or with 100 µM UDP incubation for up to 2 hours. One-Way ANOVA with post-hoc comparison to the start of UDP incubation: P≥0.8934. Scale bar, 10 µm. (solid line: mean ± SEM; dashed line: one of three wells containing 300-500 cells). (J-L) Characterization of UDP1.0 in acute brain slice. (J) Schematic illustration of the slice experiments. An AAV2/9 expressing hSyn-UDP1.0 was injected into the cortex; 3 weeks later, acute brain slices were prepared and used for two-photon imaging. (K) UDP1.0 fluorescent responses to 250 µM UDP or 250 µM ATP. Left: example image showing the UDP1.0 response to UDP or ATP. Right: overall ΔF/F mean ± SEM response from n=4-6 trials. (L) Summary data showing the response of UDP1.0 to different UDP or ATP concentration in brain slice. UDP1.0 responses were calculated by the area under the curve shown in (K) over a 60s period. Summary data are presented as the mean ± SEM from 4-8 independent trials and N=3-4 slices.

We further evaluated UDP1.0 signal dynamics in acute brain slices (Figure 1J). To achieve expression in brain tissue, the UDP1.0 sequence was inserted downstream of the human Synapsin (hSyn) promoter for neuronal targeting and packaged in AAV2/9. After delivering the virus in cortex, we could observe robust transfection within 3 weeks (Figures 1J and 1K). Exogenous application of UDP can evoke sensor responses as low as 500 nM. UDP responses are most readily observed in a dynamic range between 10-250 µM (Figures 1K and 1L). By contrast, sensor responses to ATP at 250 µM are extremely weak (Figure 1K). Even at 1 mM concentration, ATP responses are equivalent to UDP responses in the nanomolar range (Figure 1L). Therein, acute slice studies suggest that UDP1.0 can function with high specificity and selectivity for UDP over ATP in brain tissue.

### Enhanced UDP release occurs following status epilepticus

To understand endogenous UDP dynamics in the brain, both in the basal state and in epileptogenesis, we targeted UDP1.0 to neuronal membranes in cortex (AAV2/9-hSyn-UDP1.0) and performed longitudinal 2-photon imaging (chronic window preparations; Figure 2A). Across our longitudinal study, we observe two distinct types of UDP release events, based upon their rise kinetics. “Type 2” UDP events are most common across all periods and have gradual rise kinetics, ranging from seconds to tens of seconds (Figure 2B). The slower rise of Type 2 events can be modelled under conditions of prolonged UDP release *ex vivo*, while their decay kinetics follow a single-exponential slope (Figures S3G and S3H). “Type 1” UDP events are only observed *in vivo* acutely following KA-SE (2-6 hr. post-SE, representing 40.0 ± 3.3% of events in this period; Figures 2A and 2D). These events have near-instantaneous rise kinetics, which arrive upon a peak that rapidly decays, giving Type 1 events their defining ‘sharp peak’ characteristic (Figure 2B). After the initial rapid decay, Type 1 events undergo more gradual decay, similar to Type 2 events. The overall decay of Type 1 events is therefore best fit by a two-component exponential equation (Figure S3H). The near-linear rise slope of Type 1 events can be modelled by highly transient UDP release (Figure 2C). However, we cannot recapitulate the initial rapid decay of these events, suggesting a highly unique release or degradation characteristic may occur during excitotoxicity.

**Figure 2:**
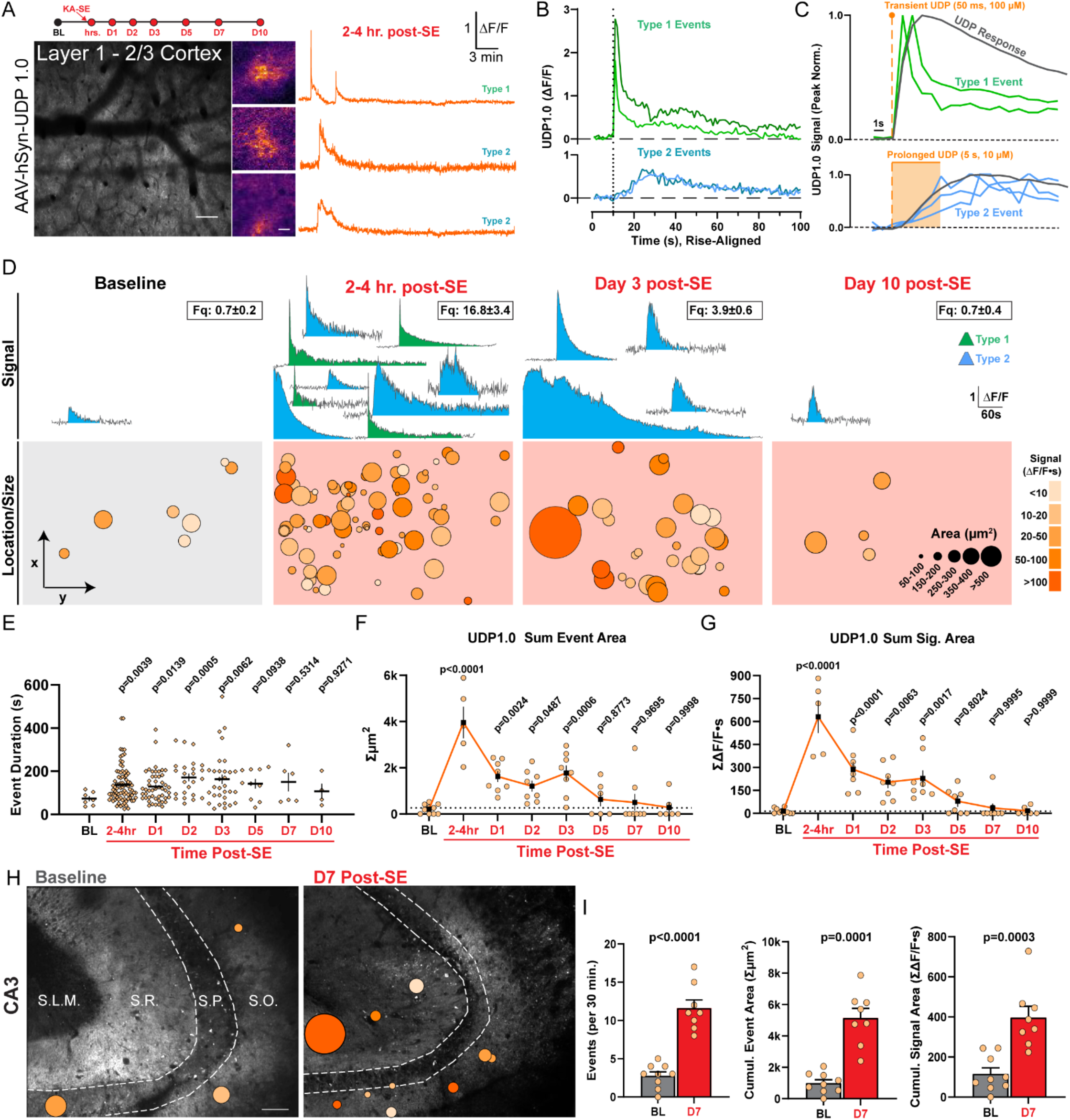
Enhanced UDP release occurs following status epilepticus. *(A) In vivo* 2-photon study timeline and image of UDP1.0 expression. Scale bar, 100µm. Examples of UDP sensor event fluorescence and corresponding ΔF/F traces. Scale bar, 20µm. (B) Examples of Type 1 sharp-peak events and Type 2 events with slower rise (ΔF/F traces aligned to rise start time). (C) Modeling Type 1 and Type 2 rise kinetics (peak normalized) against transient or prolonged UDP application *ex vivo* (gray trace). (D) Example UDP sensor signals (top, ΔF/F) and sensor event size/location under the cranial window (bottom) at baseline and during epileptogenesis. The number of ΔF/F signals displayed (top) is meant to convey frequency (frequency box represents mean ± S.E.M events per 30 min recording). Event size/location displays all events aggregated across a longitudinal cohort of N=3-5. (E) Duration of UDP release events (dot: a single event). (F) The cumulative area covered by UDP sensor events over a 30 min period (dot: one imaging region). (G) The cumulative UDP ΔF/F·s signal recorded over a 30 min period (dot: one imaging region). (H) Overlay of UDP sensor events *ex vivo* in the CA3 region of hippocampus. Scale bar, 70µm. Event scale as in (D). (I) Quantification of either UDP event frequency, cumulative UDP release area, or cumulative UDP signal area per 30 min. Welch’s T-test (dot: one or 2-3 CA3 regions surveyed from N=4 mice per time point). Data represent the mean ± SEM. D-G: Longitudinal study of 2 non-overlapping regions from N=3-5 mice. E-G: One-Way ANOVA with Fisher’s post-hoc testing vs. baseline for statistical comparison.

At baseline, UDP release events are quite rare, representing an average of 0.7 events over 30 minutes of imaging in the naïve mouse cortex (awake imaging, 620 x 620 µm field of view). However, soon after KA-SE, UDP release events increase substantially to 16.8 ± 3.4 events per 30 min (Figure 2D and Video S1), fitting the profile of a damage-associated molecular pattern (DAMP). During epileptogenesis, UDP release events are prolonged (Figure 2E), enhancing their average signal area (Figure 2D). In addition, UDP release becomes more frequent (Figure 2D). Taken together, these dynamic shifts provide evidence that microglia could be exposed to a far greater cumulative area of UDP release in the parenchyma during epileptogenesis (Figure 2F), and a greater amount of UDP release (based upon UDP1.0 ΔF/F·s signal area; Figure 2G).

We additionally surveyed UDP release dynamics in the hippocampus, because epileptogenesis engenders substantial network changes in limbic circuitry (Figure 2H). Similar to the cortex, UDP release events are infrequent in naïve hippocampal slices (2.78 ± 0.52 events per 30 min). However, one week after KA-SE, UDP release events increase 3-fold in their frequency (11.63 ± 1.07 events per 30 min; Figure 2I) with a ∼160% increase in average physical size (306 ± 29 µm^2^ naïve vs. 497 ± 87 µm^2^ D7, p=0.0194; Welch’s t-test). After epileptogenesis, there is a notable shift towards UDP release preferentially occurring in the stratum radiatum (SR) layer of both the CA3 (67%; n=49/73 total events) and CA1 regions (80% in a subset of trials; n=24/30 events). At baseline, UDP release was approximately equivalent in SR and stratum oriens (SO; 40% SO vs 44% SR). Therefore, available evidence suggests that microglia are also exposed to a greater cumulative area of UDP release and a stronger overall UDP signal during epileptogenesis in hippocampus (Figure 2I).

Finally, to determine if UDP release is a conserved response to excitotoxicity, we performed a challenge experiment in brain slices at the end of the cortex study. To induce excitotoxicity, we exchanged normal aCSF for aCSF lacking extracellular magnesium (0 Mg^2+^), which can result in epileptiform bursts within minutes^29^. Similar to KA-SE, Mg^2+^ washout induces a prominent shift in UDP signal dynamics (Figure S4A and Video S2). Type 1 “sharp-peak” UDP sensor events emerge during 0 Mg^2+^ slice conditions (17.0 ± 2.2% of events in 0 Mg^2+^; Figure S4B), which were not present at baseline. These data suggest that Type 1 events may reflect a unique form of UDP release during excitotoxicity. The prevalence of Type 1 events in 0 Mg^2+^ significantly increases the maximum amplitude and modestly increases the signal area of UDP release compared to baseline (aCSF with 1.3 mM Mg^2+^; Figure S4C). Overall, 0 Mg^2+^ excitotoxic conditions rapidly increase UDP release frequency, from an average of 6 events to 14.25 events across cortical layers (Figures S4D and S4E), providing further evidence for UDP to function as a DAMP during hyperexcitable conditions.

### Microglial calcium elevations during epileptogenesis are P2Y_6_ dependent

Enhancements in UDP release and P2Y_6_ sensitivity may significantly contribute to microglial calcium signaling during epileptogenesis. To test our hypothesis, P2Y_6_ KO (*P2ry6^-/-^*) and WT (*P2ry6^+/+^*) lines were bred to microglial calcium reporter mice (Cx3Cr1^CreER-IRES-eYFP^; R26^LSL-CAG-GCaMP6s^). *P2ry6^-/-^* microglia have a confirmed loss of *P2ry6* transcript without changes in other P2X or P2Y gene family members (Figure S5A). In slices, *P2ry6^-/-^* microglia have greatly reduced calcium responses to a specific P2Y_6_ agonist (MRS-2693) and UDP (Figures S5B-S5G), while maintaining similar calcium responses to ATP (13.02 ± 1.822 ΔF/F·s *P2ry6^+/+^* vs.12.85 ± 1.149 ΔF/F·s *P2ry6^-/-^*, Student’s T-test, p=0.9999). Relatedly, *P2ry6^-/-^* microglia have attenuated UDP chemotaxis without alterations in their ATP chemotaxis (Figures S5H and S5I). These results verify that *P2ry6^-/-^* microglia have strongly blunted UDP calcium and motility responses without adverse effects on other purine systems.

Using *in vivo* two-photon imaging (Figure 3A), we first discovered that P2Y_6_ KO has little impact on spontaneous microglial calcium signaling or process motility in a naïve state. At baseline, both populations of microglia are defined by low-to-absent spontaneous calcium signaling (WT: 81.9 ± 2.1% inactive, 15.9 ± 1.9% with low activity; KO: 84.6 ± 2.3% inactive, 13.9 ± 1.2% with low activity; Figures 3B-3D) and an equivalent frequency of process extension and process retraction (Figures 3H and 3I). Initial similarities between genotypes may reflect the rarity of UDP release in naïve cortex (Figure 2), providing little influence on P2Y_6_ dynamics at rest. On the other hand, there are substantial differences in microglial calcium signaling between genotypes during epileptogenesis. Over the first week of epileptogenesis, approximately half of all *P2ry6^+/+^*microglia shift from an inactive to a highly active calcium signaling profile (highly active: 1.9% ± 0.4% at baseline vs. 49.1 ± 3.5% at Day 7; Figures 3B-3D). However, in *P2ry6^-/-^*microglia, the transition to high-level calcium signaling is strongly attenuated (highly active: 1.8% ± 0.9% at baseline vs. 13.5 ± 2.2% at Day 7), with the majority of processes retaining low-to-absent calcium signaling profiles during early epileptogenesis (Figures 3B-3D and Video S3). Importantly, P2Y_6_ KO mice were derived and maintained from an intercross to limit any potential effects of background genetics on kainate sensitivity (aggregate dose to reach a first seizure: 25.74 ± 1.39 mg/kg KA, *P2ry6^+/+^*; 25.78 ± 1.18 mg/kg KA, *P2ry6^-/-^*; Student’s T-test, P=0.9801). Further, mice were only enrolled in studies if they met strict seizure severity inclusion criteria (see Star Methods). We would therefore conclude that microglia without P2Y_6_ receptors have substantially limited calcium signaling in the first two weeks of epileptogenesis (Figure 3E).

**Figure 3:**
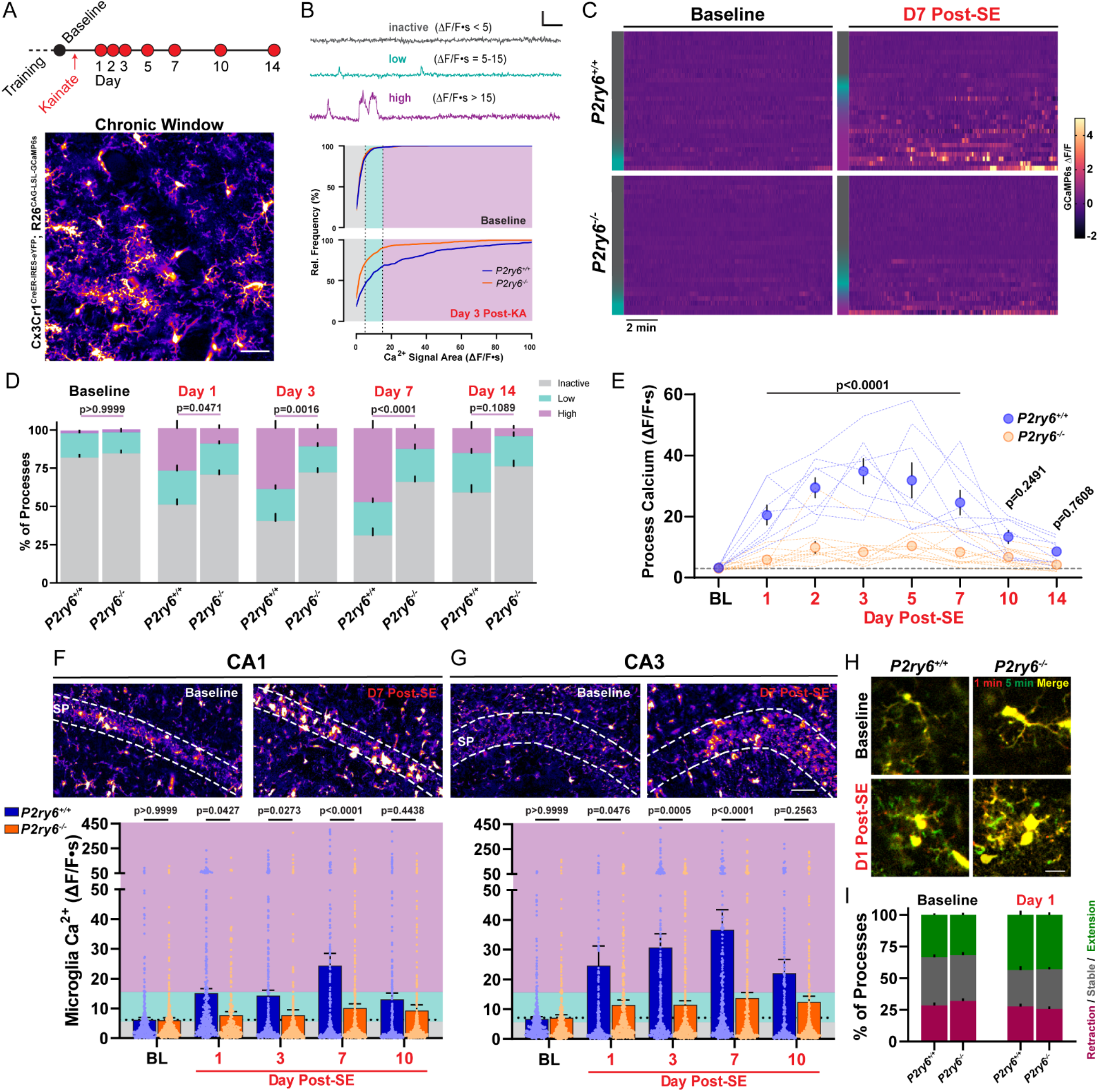
Microglial calcium elevations during epileptogenesis are P2Y6 dependent. (A) Experimental timeline for *in vivo*, longitudinal calcium imaging of cortical microglia. Representative field of view. Scale bar, 40µm. (B) (Top) Examples of inactive, low, and high-level microglial calcium signaling used for segmenting activity. Scale bar: 1-fold ΔF/F and 60s. (Bottom) Microglial calcium activity distributions (ΔF/F·s) at baseline and 3 days after KA-SE. (C) Heatmaps of microglial GCaMP6s calcium activity (ΔF/F) by genotype and time period (30 ROIs chosen to match inactive/low/high activity distributions—left color bar). (D) Percentage of total microglia ROIs exhibiting no, low, or high activity between genotypes and across periods of epileptogenesis. Two-way ANOVA with Sidak’s post-hoc test between activity levels (P-value compares high-level activity). (E) Overall calcium activity in *P2ry6^+/+^*and *P2ry6^-/-^* microglia across two weeks of epileptogenesis. Two-way ANOVA with Sidak’s post-hoc test (dotted lines represent a longitudinal imaging region; dots represent group mean ± SEM). (F) (Top) 2P images of microglia in the CA1 region from acute brain slice. (Bottom) Spontaneous calcium activity in CA1 microglia across epileptogenesis. Two-Way ANOVA with Sidak’s post-hoc test (dots represent one ROI; survey of 2 slices per mouse; N=4 mice per time point and genotype). (G) As in (F), for microglia of the CA3 region. Scale bar, 50 µm. (H) *In vivo* average intensity images of microglia (overlay of start vs. 5 min) to indicate process motility. Scale bar, 15µm. (I) Overall percentage of microglial processes that extend outward, remain stable, or retract during *in vivo* imaging. Two-way ANOVA with Sidak’s post-hoc test: no significant motility differences between genotypes (p=0.5505-0.9965 by motility type). Bars represent the mean ± SEM. *In vivo* (A-E): N=4 *P2ry6^+/+^*mice, N=6 *P2ry6^-/-^* mice with 2 non-overlapping regions surveyed per mouse.

In acute brain slices, we also evaluated how P2Y_6_ KO impacts microglial calcium signaling in hippocampus. As an important comparative control, cortical microglia display reasonably similar patterns of calcium activity in both slices and *in vivo* (Figures S6A-S6C). At baseline, *P2ry6^+/+^* and *P2ry6^-/-^* microglia rarely exhibit spontaneous calcium activity in hippocampus (Figures 3F, 3G, and S6D). However, *P2ry6^+/+^* microglia begin to display clear enhancements in spontaneous calcium activity 1, 3, 7, and 10 days after KA-SE, while *P2ry6^-/-^* microglia have significantly attenuated spontaneous calcium signaling in hippocampus throughout early epileptogenesis (Figures 3F and 3G). Taken together, we conclude that enhanced spontaneous microglial calcium activity is a wide-spread feature of epileptogenesis and is primarily governed by P2Y_6_ signaling in microglia.

### P2Y_6_ activation regulates lysosome expression and facilitates neuronal soma engulfment

The function of P2Y_6_ signaling in epileptogenesis is not definitively known. However, P2Y_6_ activation can promote phagocytosis in both sterile^25,30-33^ and infection-based contexts^25^. We therefore investigated how P2Y_6_ signaling influences phagocytosis during epileptogenesis. In WT mice, we see a progressive expansion of CD68 phagolysosome area (example CA3 time course: Figures S7A and S7B), with co-staining confirming expression in IBA1 cells. Within 3-7 days after KA-SE, *P2ry6^+/+^* microglia robustly up-regulate lysosomes in the hippocampus, amygdala, and thalamus, while *P2ry6^-/-^*microglia have substantially reduced lysosome expression in these regions (Figures 4A and 4B). Importantly, the hippocampus, amygdala, and thalamus are all limbic regions known to have a high concentration of dead or dying neurons following systemic KA-SE^8,9^. As an important control, baseline levels of CD68 expression are equivalently low in the naïve state for both genotypes (Figure S7B). Surprisingly though, P2Y_6_ KO does not influence morphology in microglia, as the progression and extent of ameboid transition is similar between genotypes in hippocampus (Figures 4C and 4D). Further evaluation suggests that initial seizure severity may strongly predict the extent of ameboid transition in microglia, independent of genotype. On the other hand, the *P2ry6^-/-^* genotype substantially limits the extent to which seizure severity produces greater lysosome expression (Figure 4E). Therefore, P2Y_6_ signaling has the unique ability to uncouple lysosome up-regulation from an ameboid (classically “phagocytic”) state transition.

**Figure 4:**
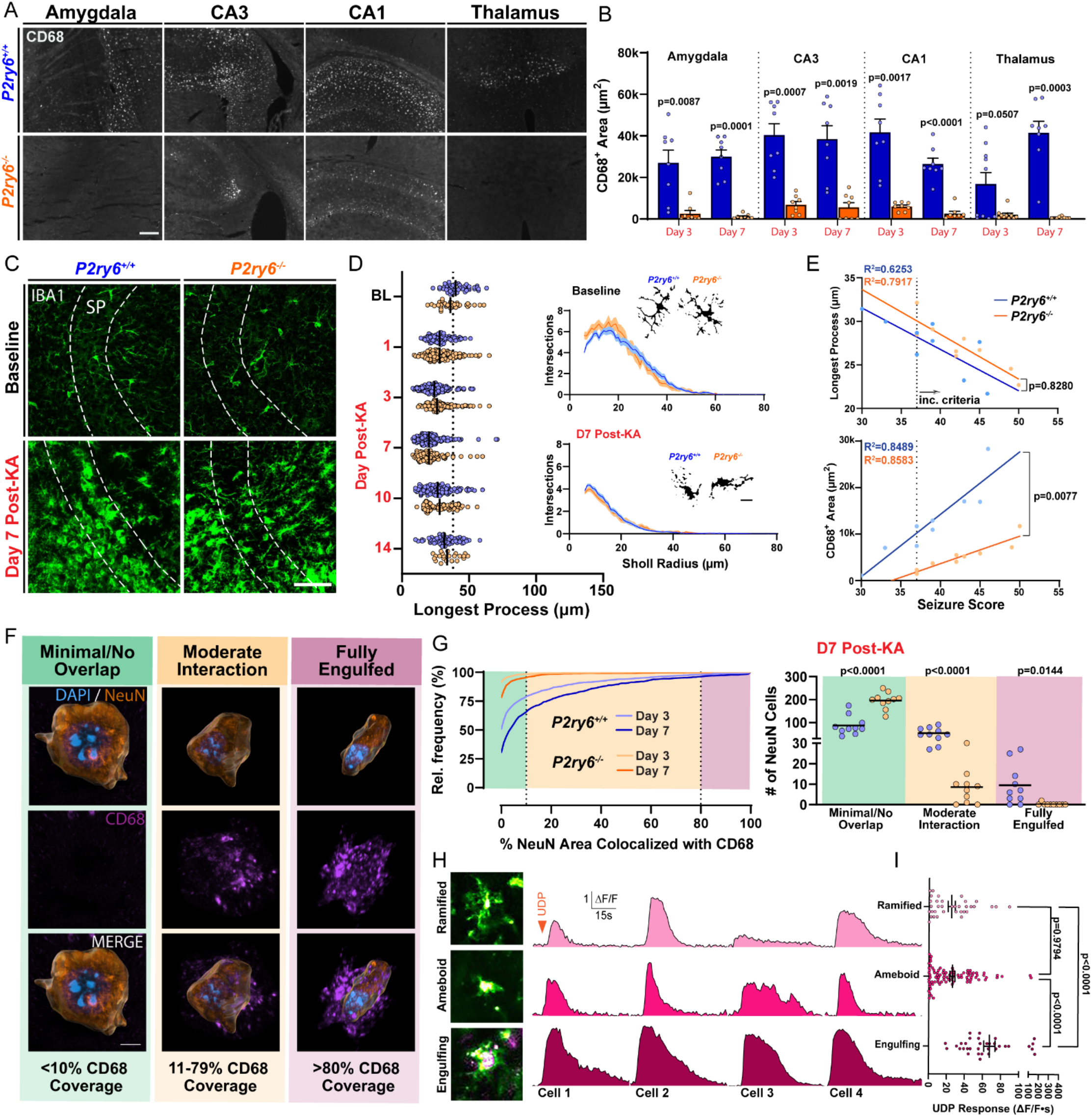
P2Y6 activation regulates lysosome expression and facilitates neuronal soma engulfment. (A) CD68 lysosome expression across limbic regions 3 days after KA-SE. Scale bar, 100 µm. (B) Quantification of CD68 area by region and genotype 3- and 7-days after KA-SE. One-way ANOVA with Dunnett’s post-hoc test by region (dots: one region; two regions from N=4-5 mice/group). (C) IBA1 staining in the CA3 region by genotype and time point. Scale bar, 40µm. (D) (Left) CA3 microglia longest process analyses. Two-Way ANOVA effect of genotype: F_(1, 1094)_=0.5979, p=0.4395 (dot: one cell). (Right) Sholl plots with representative cell morphologies. Scale bar, 5µm. Survey of 10-40 microglia per mouse; N=3-5 mice/group. (E) Correlations between initial seizure severity and CA3 microglia process length or CD68 area. Simple linear regression (dot: one mouse; aggregated from day 3 and day 7 time points). (F) Histological classification of NeuN neuron (Imaris surface rendering) and CD68 phagolysosome interactions in CA3 from 3D confocal microscopy, ranging from no/minimal interaction to full engulfment. Scale bar, 5µm. (G) (Left) Distribution of NeuN neurons based upon their CD68 coverage (survey of n=1500-1989 CA3 neurons from 10 CA3 subfields and N=5 mice/group). (Right) The number of CA3 neurons having different CD68 coverage. One-Way ANOVA with Sidak’s post-hoc test (dots: one CA3 subfield). (H) Representative images of ramified, ameboid, and neuron-engulfing microglia in CA3 brain slice (purple: transient intercalation of AF568 dye with a cell soma). Representative ΔF/F calcium responses to 500 µM UDP. (I) Comparison of CA3 microglia UDP calcium responses by morphology. One-way ANOVA with Tukey’s post-hoc test (dot: one microglial cell; survey of cells from WT brain slices prepared 3, 7, and 10 days post-SE).

In regions such as CA3, CD68 lysosome upregulation was most substantial in the pyramidal cell layer (SP; Figure S7C), which contains the soma of principle neurons. We therefore evaluated whether microglia were performing phagocytosis of neuronal somata (Figure 4F and Video S4). CD68 (Lysosome)–NeuN (Neuron) interactions were qualified as either no/minimal coverage (CD68 area encompasses <10% of the NeuN soma), moderate coverage (CD68 area encompasses 11-79% of the NeuN soma), or full engulfment (CD68 area encompasses >80% of the NeuN soma). At the peak of CD68/NeuN interactions in *P2ry6^+/+^* tissue (day 7 after KA-SE), approximately 40% of all CA3 neurons surveyed had a moderate level of interaction with CD68 microglia (54 ± 7 pyramidal neurons per 212 x 212 x 20µm volume; Figure 4G). At this peak time point, we also observed approximately 10 neurons per imaging field reaching full engulfment criteria (6.3%, or n=95 of 1501 neurons; Figure 4G). In *P2ry6^-/-^* mice, virtually no CA3 neurons undergo full engulfment (0.13%, or n=2 of 1529 neurons; Figure 4G). Interactions between CD68 and NeuN were chosen as a primary metric, because they could more directly suggest phagocytic outcomes. However, in performing the same analysis using IBA1 as a marker, we similarly observe that IBA1 microglia are less likely to interact with and engulf CA3 neurons in *P2ry6^-/-^* hippocampus (Figures S7F and S7G). Notably, *P2ry6^-/-^* IBA1 cells appear to have a deficit in either their ability to migrate into the pyramidal layer or expand therein, based upon IBA1 signal changes over time (Figures S7D and S7E). These results suggest that *P2ry6^-/-^* microglia have a severely limited ability to perform somatic phagocytosis in epileptogenesis.

Additional observations also suggest a potential link between calcium activity and the phagocytotic process. First, microglia in the CA3 region have the highest spontaneous calcium activity of any area studied (Figure S6D), which peaks at the height of phagocytic engagement—day 7 post-SE (Figure 3G). Second, in surveying UDP-based calcium responses, we observed that AlexaFluor dye application can transiently intercalate into neuronal cell membranes. As a result, we could identify microglial-neuron interactions and their associated UDP calcium responses. Microglia potentially engulfing neurons in brain slice had particularly large UDP calcium responses in CA3 (Figure 4H). These microglia exhibit UDP calcium responses that are approximately 3x larger than either ramified or ameboid microglia in the same imaging field (Figure 4I). Therein, we observe multiple structural and functional relationships between P2Y_6_ calcium signaling and phagocytosis: microglia with the strongest UDP calcium responses appear to perform phagocytosis, while microglia without the P2Y_6_ pathway are severely limited in their ability to up-regulate lysosomes and fully engulf neurons.

### Attenuating calcium signaling through CalEx phenocopies lysosome impairments in microglia

To pinpoint which aspects of phagocytosis may specifically be related to the downstream calcium activity of the P2Y_6_ pathway, we employed a parallel approach to reduce calcium signaling. ATP2B2 is a transmembrane protein that facilitates the rapid removal of calcium from the cytosol, reducing its signaling time^34^ (Figure 5A). ATP2B2 can be expressed in a Cre-dependent manner through the Calcium Extruder (“CalEx”) mouse line^35^ (R26^LSL-CAG-ATP2B2-mCherry^). To achieve high microglial specificity, we bred the CalEx line to TMEM119^2A-CreERT2^ mice. Using the mCherry tag expressed in the CalEx system, we confirmed that CalEx recombination occurs selectively in microglia (<1% of recombination outside of IBA1^+^ cells) and in a Cre-dependent manner (<1% recombination in the absence of tamoxifen; Figure 5B). However, in naïve mice, recombination was not robust across brain regions (20.9-27.2% mean recombination across amygdala, cortex, CA1, and CA3; Figure 5C), which precludes intensive study at the systems level. In a second cohort used for epileptogenesis studies, we increased tamoxifen administration and could observe more robust recombination in hippocampal microglia (mean recombination: 42.0% in CA3 and 41.2% in CA1; Figure 5C). To validate CalEx function, we used an improved viral delivery system to target GCaMP6f to CA1 microglia^36^ (rAAV-SFFV-DIO-GCaMP6f-WPRE-hGH) and performed calcium imaging *ex vivo* (Figure 5D). In GCaMP6f-expressing control microglia (TMEM119^2A-CreERT2^; R26^wt/wt^), focal UDP application leads to an expected, long-lasting calcium response in slices (42.0 ± 1.6s; Figures 5E-G). However, in microglia with confirmed mCherry-CalEx expression (Figure 5E), UDP application results in a significantly shortened calcium response (16.8 ± 1.3s; Figures 5F and 5G), demonstrating that CalEx can reduce the time course of purine-driven microglial calcium elevations.

**Figure 5:**
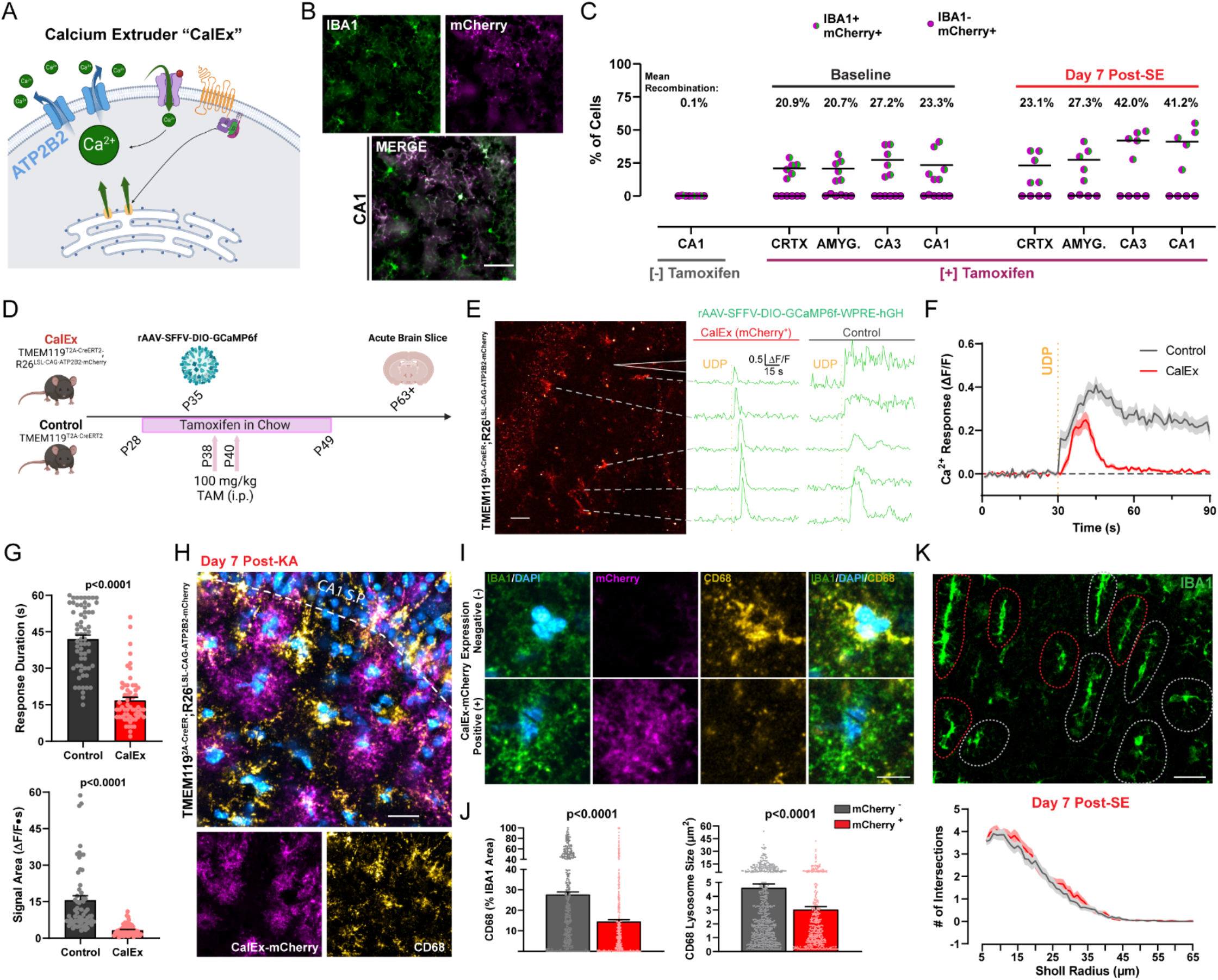
Attenuating calcium signaling through CalEx phenocopies lysosome impairments in microglia. (A) Illustration of ATP2B2 calcium extruder (“CalEx”) function. (B) Images of CalEx-mCherry recombination against an IBA1 co-stain in TMEM119^2A-CreER^;R26^LSL-CAG-ATP2B2-mCherry^ mice treated with tamoxifen. Scale bar, 30 µm. (C) Quantification of mean recombination by region (dot: one mouse; N=5-6 mice per group). (D) Outline of experimental steps to validate CalEx function in microglia. (E) Example of mCherry fluorescence in an acute brain slice with corresponding ΔF/F calcium responses following 500 µM focal UDP application (“CalEx” tissue). Scale bar, 40 µm. Right ΔF/F traces: example UDP calcium responses in an acute slice prepared from the control mouse line. (F) Overall calcium response (ΔF/F) to UDP application (lines represent the mean ± SEM; aggregate of n=69 responding cells in Control tissue and n=59 responding cells in CalEx tissue; survey of 2-4 slices from N=4-5 mice per group). (G) Quantification of 500 µM UDP calcium response duration, and signal area between Control and CalEx microglia. Student’s T-test (dot: one cell; n=69 control cells and n=59 CalEx-mCherry cells; 2-4 slices from N=4-5 mice/group). (H) Representative image of CalEx-mCherry and CD68 expression in CA1 SR one week after KA-SE. Scale bar, 25 µm. (I) Closer evaluation of IBA1 and CD68 expression in a representative mCherry^+^ and mCherry^-^ cell from (D). Scale bar, 7 µm. (J) Quantification of average lysosome size and overall CD68 area (normalized to IBA1 area) for mCherry^+^ and mCherry^-^ cells. Student’s T-test (dot: one cell; survey of n=412 mCherry^+^ microglia and n=568 mCherry^-^ microglia in CA1 SR from N=5 mice/group). (K) IBA1 morphology in CA1 SR with red and gray outlines denoting mCherry^+^ and mCherry^-^ cells, established from the mCherry channel. Scale bar, 20 µm. Sholl plots for mCherry^+^ and mCherry^-^ cells one week after KA-SE (n=88 mCherry^+^ and n=103 mCherry^-^ cells from N=5 mice/group). Bar or line graphs display the mean ± SEM.

In the process of characterizing CalEx function, we observed an interesting patchwork of CD68 expression in CA1 SR one week after SE (Figure 5H). Upon further investigation, we found that IBA1 cells with CalEx expression (mCherry^+^) had significantly reduced CD68 levels compared to adjacent mCherry^-^ IBA1 cells in the same tissue (Figure 5I). In mCherry^+^ microglia, CD68 lysosomes were smaller in size and occupied less IBA1 cell area (Figure 5J), suggesting that calcium extrusion may also prevent lysosome maturation and expansion in hippocampal microglia. Interestingly, adjacent mCherry^+^ and mCherry^-^ microglia do not differ in their IBA1-based Sholl morphologies, despite differences in CD68 expression (Figure 5K). Thus within-subject, cell-based analyses of CalEx microglia further support the importance of calcium signaling for microglial lysosome regulation, without clear impact on ameboid transition.

### Enhanced pro-inflammatory signaling is coordinated with phagocytosis in *P2ry6^+/+^* microglia

We additionally turned to bulk RNA-seq analyses to potentially uncover unique roles for the P2Y_6_ pathway in regulating microglial function. Microglia were isolated from whole brains of *P2ry6^+/+^* and *P2ry6^-/-^* mice three days after KA-SE (Figure 6A), which corresponds to the peak period of microglial spontaneous calcium signaling in the cortex (Figure 2E). In comparing transcriptomes between genotypes soon after KA-SE, a highly specific profile emerged, with most differentially expressed genes (DEGs) mapping on to immune-related function (Figure 6B). In particular, P2Y_6_ function may be closely tied to *Trim* gene regulation, as this gene family represents 7 of the top 30 DEGs. Tripartite motif-containing (TRIM) genes encode a superfamily of over 60 E3 ubiquitin ligases, which are classically known for their role in innate immune activation following infection^37^. TRIM genes also regulate multiple inflammatory pathways, including Type I interferon (IFN) and NF-κB transcription^38,39^, reflected in many of enriched Gene Ontology (GO) processes (Figure 6C). Even though SE represents a sterile insult, there are key points of overlap between the immune profiles seen in response to seizure damage and infection, as noted by other epilepsy researchers^40,41^. Critically, top upregulated genes in WT mice, *Trim12* (human Trim5 ortholog), *Trim 30b*, *Trim30d* (human Trim79 ortholog), and *Trim34a* are all positive regulators of NF-κB transcription, while top down-regulated genes, *Mturn* and *Trim21*, are negative regulators of NF-κB transcription^37,38^. From these insights, we further evaluated P2Y_6_ signaling through an immunological lens, by assessing its links to NF-κB-related inflammatory pathways.

**Figure 6:**
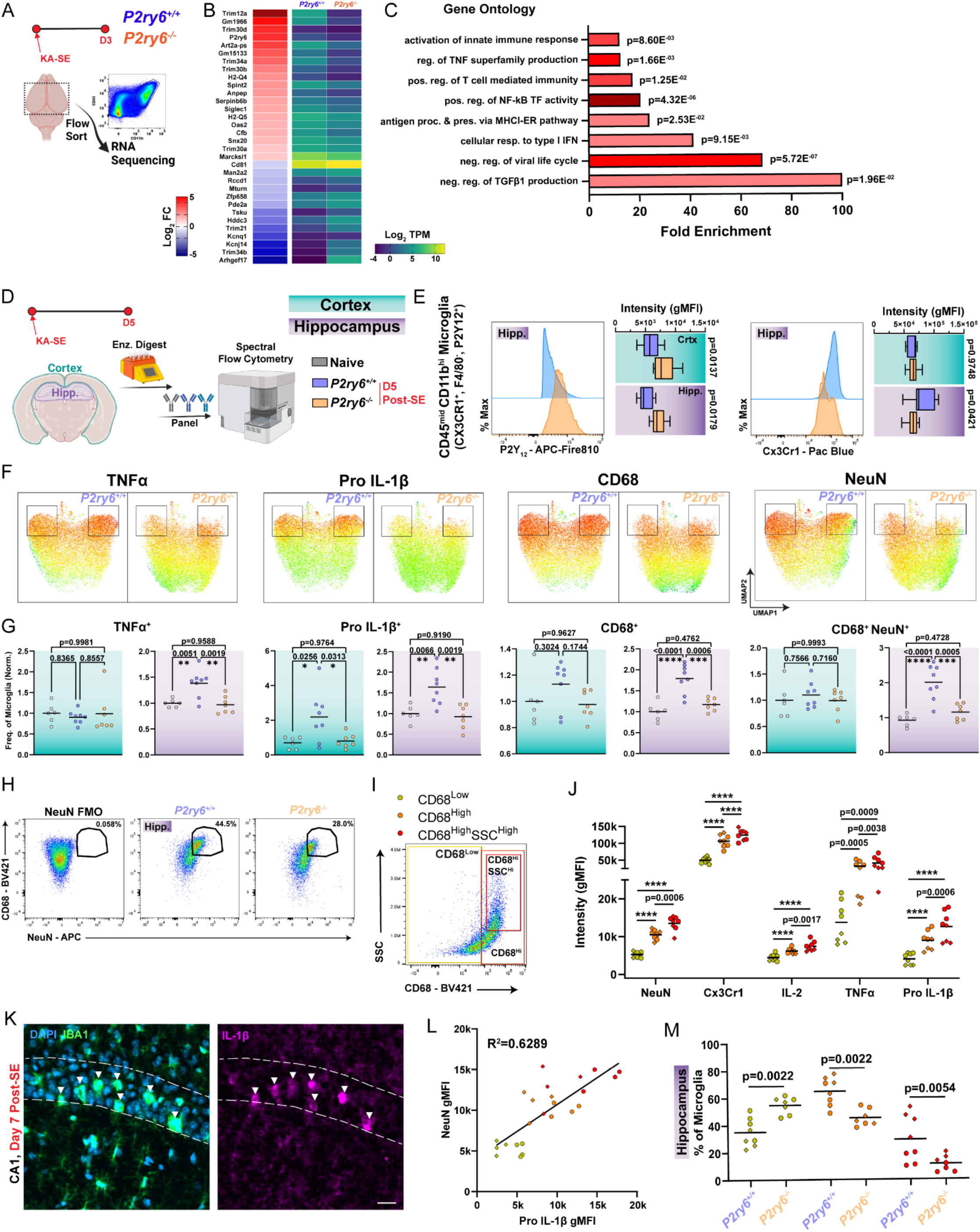
Enhanced pro-inflammatory signaling is coordinated with phagocytosis in *P2ry6^+/+^* microglia. (A) Outline of transcriptomics experiment using FACS-based microglia isolation 3 days after KA-SE from whole brain. (B) Top differentially expressed genes (DEGs) in microglia between WT-KO genotypes (log_2_FC>1.0 and BH-adjusted p<0.1). (C) Key Gene Ontology (GO) terms based upon the DEG set in (B). Details are provided in the Star Methods section. (D) Protein-level evaluation of hippocampal and cortical microglia using high-parameter flow cytometry 5 days after KA-SE. (E) Mode-normalized, representative histograms and intensity distributions for microglial P2Y_12_ and Cx3Cr1 expression. Two-Way ANOVA with Sidak’s post-hoc test (box plots display the mean and interquartile range with whiskers denoting min to max). (F) UMAPs of TNFα, Pro IL-1β, and CD68 expression, or NeuN/CD68 co-detection in hippocampal microglia (boxes highlight a distinct population between genotypes). (G) Frequency of marker expression by microglia in cortex and hippocampus. One-Way ANOVA with Tukey’s post-hoc test (normalized to mean naïve frequency per cohort/batch; dot: one mouse). (H) Pseudocolor plots displaying the NeuN gate, based upon an FMO, and application of that gate onto hippocampal microglia. (I) Psuedocolor plot with gates to isolate microglial populations which are CD68^Low^, CD68^High^, and CD68^High^ SSC^High^. (J) Evaluation of key signal intensity differences between *P2ry6^+/+^* hippocampal microglia which are CD68^Low^, CD68^High^, and CD68^High^ SSC^High^. One-Way ANOVA with Tukey’s post-hoc test (dot: one mouse, where dot shape denotes cohort; ****p<0.0001). (K) Representative immunofluorescent imaging of DAPI, IBA1, and IL-1β expression in WT CA1 one week after KA-SE. Scale bar, 20µm. (L) Correlations between Pro IL-1β intensity and NeuN intensity inside of microglia. Simple linear regression (dot: one mouse). (M) Comparison of different CD68 population frequency between genotypes in the hippocampus. Two-Way ANOVA with Sidak’s post-hoc test (dot: one mouse, where dot shape denotes cohort). Data were aggregated from N=6 naïve mice (4 WT, 2 KO) N=8 D5 *P2ry6^+/+^* mice and N=7 D5 *P2ry6^-/-^* mice across two independent cohorts.

We used high-parameter flow cytometry to analyze microglia and their activation states 5 days after KA-SE in hippocampus and cortex (Figures 6D and 6E). A day 5 time point should reasonably match day 3 transcriptional analyses, while capturing aspects of both innate and early adaptive immunity (Figure 4). NF-κB activation can result in the transcription of multiple cytokines, including IL-1, IL-6, and TNFα. In UMAP plots, we observe that a subset of *P2ry6^+/+^* hippocampal microglia have enriched TNFα and Pro IL-1β expression (Figure 6F). Further evaluation shows that *P2ry6^+/+^* microglia more frequently express TNFα than naïve or *P2ry6^-/-^* populations in the hippocampus (Figure 6G). Shifts in Pro IL-1β expression are also highly pronounced, with *P2ry6^+/+^* microglia more frequently expressing this cytokine in both cortex and hippocampus compared to naïve or *P2ry6^-/-^* populations (Figure 6G). In addition, Pro IL-1β gMFI is slightly increased in hippocampal *P2ry6^+/+^* microglia. Taken together, transcriptomics and flow cytometry findings support a potential role for P2Y_6_ signaling in driving NF-κB related inflammation.

We additionally evaluated differences in phagocytic populations between *P2ry6^+/+^* and *P2ry6^-/-^* microglia using flow cytometry. Consistent with immunofluorescence staining, we find that *P2ry6^+/+^* microglia in hippocampus more frequently and more robustly express CD68 phagolysosomes (Figures 6F and 6G). After gating for microglial populations (Figures S8A and S8B), we evaluated the extent of NeuN signals in microglia between genotypes. Microglia from the *P2ry6^+/+^*hippocampus are much more likely to have a co-detected NeuN signal than *P2ry6^-/-^*microglia, which we interpret as greater neuronal phagocytosis in early epileptogenesis (Figures 6F-6H). Interestingly, in UMAP visualizations, we observe that the same populations of TNFα^High^ and Pro IL-1β^High^ *P2ry6^+/+^* microglia are also most likely to express high levels of CD68 and include NeuN signals (Figure 6F). With flow cytometry, we had the ability to identify and analyze unique populations of CD68 microglia, including microglia with low CD68 expression, high CD68 expression, and high CD68 expression in combination with high side-scatter (SSC)—an indication of greater cell density (Figure 6I). In WT microglia, the intensity of NeuN signal progressively increases between CD68^Low^, CD68^High^, and CD68^High^-SSC^High^ populations, suggesting they may perform more neuronal phagocytosis (Figure 6J). Each population is also increasingly more activated based upon its Cx3Cr1 intensity (Figure 6J). As expected from UMAP visualizations, both CD68^High^, and CD68^High^-SSC^High^ microglial populations have substantially greater TNFα expression than CD68^Low^ populations, while IL-2 and Pro IL-1β expression increase significantly between CD68^Low^, CD68^High^, and CD68^High^-SSC^High^ populations (Figure 6J). Taken together, these observations suggest that pro-phagocytotic microglia are also more likely to be strongly pro-inflammatory.

Existing lines of evidence also suggest that IL-1β may play a key role in the process of neurodegeneration^42^, including cell loss during epileptogenesis^43^. Towards this end, spatial information provided by immunofluorescent staining reveals that IBA1 microglia expressing the highest levels of IL-1β may be actively phagocytosing neurons based upon their pyramidal band location and morphology (Figure 6K). Additionally, we observe strong positive correlations between NeuN signals in microglia and their Pro IL-1β expression levels in flow cytometry (Fig. 6L), supporting a potential relationship between IL-1β and neurodegeneration.

The microglial P2Y_6_ pathway may enhance hippocampal inflammation through two major routes. First, pro-inflammatory, pro-phagocytic microglia are much more prevalent in *P2ry6^+/+^* hippocampus. They do not substantially emerge in microglia without the P2Y_6_ receptor (Figures 6F and 6M). Second, the *P2ry6^+/+^* hippocampus experiences more CD45^High^ immune cell infiltration (Figure S8C). Specifically, more myeloid cells are found in the *P2ry6^+/+^* hippocampus 5 days after KA-SE (range: 0.8-19.9% of live cells in *P2ry6^+/+^* vs. 0.2-2.69% of live cells in *P2ry6^-/-^*; Figure S8D). Infiltrating myeloid cells are highly pro-inflammatory, having greater Pro IL-1β expression than even activated microglia (Figure S8E). Interestingly, the myeloid populations found in *P2ry6^+/+^* hippocampus are more intensely and more frequently MHCII^+^ than the populations found in *P2ry6^-/-^* hippocampus (Figure S8E), which may have later implications for antigen presentation and modulation of CD4 T-cell responses. Ultimately, multiple mechanisms primary and secondary to microglial P2Y_6_ function enhance the inflammatory status of the hippocampus.

### P2Y_6_ KO mice have improved CA3 neuronal survival and enhanced cognitive task performance

Enhanced hippocampal inflammation may explain why we observe indications of poorer neuronal health in *P2ry6^+/+^* hippocampus. In flow cytometry, neurons in the *P2ry6^+/+^* hippocampus are twice as likely to express TNFα 5 days after seizures (Figure S8G), while astrocyte TNFα expression remains unchanged between genotypes (Figure S8F). Histological evaluation (FluoroJade-C; FJC) suggests neuron death or damage is initially similar between genotypes 1 and 3 days after KA-SE in the amygdala, hilus, CA3, and CA1 regions (Figures S9A and S9B). Therefore, neither genotype appears to be resistant to limbic cell damage. Both genotypes also display similarly high levels of immediate early gene expression one day after KA-SE (c-Fos; Figures S9C and S9D)^44,45^. However, FJC staining levels begin to separate between genotypes one week after KA-SE. In the amygdala and CA3, FJC levels start off high in both genotypes (within 24 hrs.), but recede significantly by Day 7 in KO tissue while remaining elevated in WT tissue (Figures S9A and S9B). In the WT CA1 region, FJC positivity increases over one week, but this progression does not occur in P2Y_6_ KO mice (Figures S9A and S9B). These observations and their temporal alignment with flow cytometry findings may suggest an important role for the P2Y_6_ pathway in influencing cytotoxicity.

As a potential consequence of attenuated phagocytic interactions, inflammation, and immune infiltration, we observe greater CA3 neuron survival in KO tissue (Figures 7A-7C). In *P2ry6^+/+^* mice, loss of NeuN in the CA3 pyramidal band progressively increases after KA-SE for a one-week period, matching FJC observations (Figures 7B and 7C). Compared to baseline, approximately 40% of neurons may be lost in the CA3 pyramidal band (baseline: 13.55±0.82 neurons/100µm^2^, Day 14: 8.51±0.42 neurons/100µm^2^). However, in *P2ry6^-/-^* mice, NeuN loss does not appear to progress beyond 24 hours after KA-SE, with a consistent observation that only 15% of CA3 neurons are lost across histological time points (baseline: 13.19±0.31 neurons/100µm^2^, Day 14: 11.62±0.67 neurons/100µm^2^; Figures 7B and 7C). These findings are also corroborated by Cresyl Violet staining in Day 10 and Day 14 tissue (Figures S9E and S9F). We next assessed whether differences in observed CA3 neuron loss had a significant impact on cognition.

**Figure 7:**
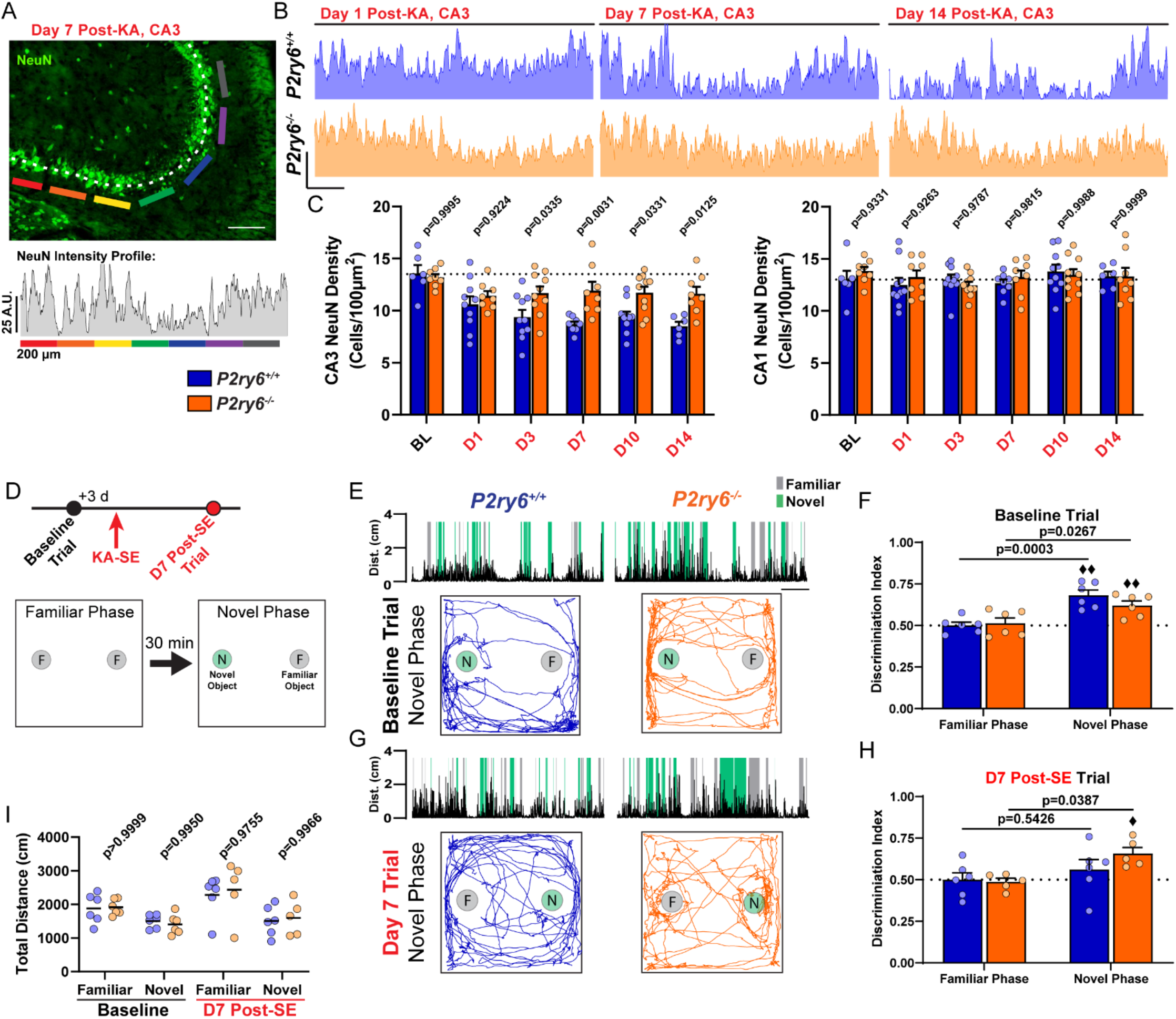
P2Y_6_ KO mice have improved CA3 neuronal survival and enhanced cognitive task performance. (A) Example of a NeuN intensity plot from day 7 post-SE WT tissue. Scale bar, 200 µm. (B) Representative NeuN intensity profiles from the CA3 region of *P2ry6^+/+^* and *P2ry6^-/-^* tissue. Scale bar, 25 A.U. and 200 µm. (C) Quantification of NeuN cell density in the CA3 or CA1 pyramidal band. Two-way ANOVA with Sidak’s post-hoc test (dots: one region; bilateral survey from N=3-5 mice/group). (D) Timeline of Novel Object Recognition (NOR) task (repeated at baseline and day 7 post-SE). Outline of the task procedure. (E) Representative trial performance in the novel phase during the baseline (naïve) test. (Top) Distance moved with periods highlighted during novel or familiar object interaction. Scale bar, 1 min. (Bottom) Trace of mouse movement in the arena. (F) Discrimination index score by genotype and trial phase during the baseline (naïve) test. Two-Way ANOVA with Sidak’s post-hoc test between familiar and novel phase performance by genotype (dot: one mouse; ♦♦P<0.01, one-sample T-test comparison to chance: 0.50). (G) As in (E), representative trial performance 7 days after KA-SE. (H) As in (F), discrimination index 7 days after KA-SE. Two-Way ANOVA with Sidak’s post-hoc test between familiar and novel phase performance by genotype (dot: one mouse; ♦P<0.05, one-sample T-test comparison to chance). (I) Total distance moved by trial phase. Two-Way ANOVA with Sidak’s post-hoc test (dot: one mouse). *P2ry6^+/+^* and *P2ry6^-/-^* mice used longitudinally at baseline and day 7 post-SE (N=6 mice/group at baseline; N=6/6 surviving *P2ry6^+/+^*; N=5/6 surviving *P2ry6^-/-^* at day 7 post-SE).

We assessed cognitive performance in the novel object recognition (NOR) task, known to rely on CA3 function in associating objects and spatial locations^46,47^. NOR performance was assessed in the same cohort of *P2ry6^+/+^* and *P2ry6^-/-^* mice at baseline (naïve state) and one week following KA-SE (Figure 7D). In the baseline test, naïve mice first explored two identical objects in the familiarization phase, with an expected lack of object preference seen in both genotypes (50:50 ratio; Figure 7F). In the novel phase (30 min delay), both *P2ry6^+/+^* and *P2ry6^-/-^* mice preferentially explored the novel object over the familiar object, resulting in a discrimination index score that was above both chance (0.50; one-sample T-test) and their discrimination scores in the familiar phase (Figures 7E and 7F). One week after KA-SE (10 total days after the prior test), we re-tested mice in NOR using different object sets. Familiarization phase performance occurred at the expected 50:50 ratio (Figure 7H). However, in the novel object phase, *P2ry6^+/+^* mice failed to explore the novel object at levels that were significantly distinguishable from chance or their performance in the familiarization phase (Figures 7G and 7H). On the other hand, *P2ry6^-/-^* mice still maintained their ability to distinguish a novel object from a familiar object one week after KA-SE, based upon above-chance and above-familiar phase task performance (Figures 7G and 7H). Using an open-source animal tracking program (EZ track^48^), we found animal movement was similar between genotypes for all test periods/phases (Figure 7I), indicating animal mobility was not a confounding factor. Therefore, NOR suggests that mice without P2Y_6_ pathway activity can better accomplish cognitive tasks in early epileptogenesis, likely owing to significant reductions in network inflammation and cell loss during this period.

## Discussion

As innate immune cells of the CNS, microglia are well positioned to sense and respond to local shifts in brain homeostasis, including hyperactivity, inflammation, and cell damage. Multiple lines of evidence suggest that microglia engage calcium activity in response to these cues^3-7,49^, but the definitive pathways and downstream consequences have not been elucidated. Using a high-performance UDP biosensor, we discovered that UDP release in parenchyma commonly occurs in response to hyperexcitability and remains elevated for multiple days following KA-SE. UDP can activate calcium signaling in microglia through the P2Y_6_ pathway, which serves as a major impetus for spontaneous calcium activity during early epileptogenesis. As a ubiquitous secondary messenger, calcium signaling performs multiple roles in microglia with interesting regional dependency. In the hippocampus, attenuation of microglial calcium via P2Y_6_ KO or CalEx expression results in a failure to robustly upregulate CD68 lysosomes in response to neuronal damage. Flow cytometry additionally indicates that P2Y_6_ KO prevents pro-inflammatory cytokine production in hippocampal microglia and reduces myeloid infiltration. Therein, calcium signaling coordinates pro-inflammatory and pro-phagocytic state transitions in microglia, shaping the early neuroimmune landscape in the hippocampus. As a consequence, P2Y_6_ signaling results in enhanced neurodegeneration and poorer cognitive outcomes.

One of the key enabling factors for our research is the ability to longitudinally study UDP using a GRAB_UDP1.0_ sensor. Compared to ATP/ADP, UDP signaling is relatively understudied, due in part to issues with antibody specificity^27^, and the lack of brain-permeable, specific antagonists^24^. Biosensors provide the ability to observe UDP levels in real time, particularly when release occurs as a damage signal/DAMP in the extracellular space. To date, purine release mechanisms are debated in the CNS. In the spinal cord, ATP release can clearly occur through vesicular mechanisms following spinal cord injury^16^. In the brain, available evidence more likely implicates a volume-regulated channel^50^ or hemi-channel system^51,52^ for neuronal ATP release. Virtually no studies provide an indication of how UDP (or more likely UTP) is released. Our studies in brain slice under 0 Mg^2+^ conditions suggest that UDP release can be a rapid response to excitotoxicity. The acute excitotoxic effects of both 0 Mg^2+^ (within minutes) and kainate seizures (within minutes-hours) induces a unique form of UDP fluorescence, termed Type I or sharp peak events. In studies of the CA3, we were intrigued to see UDP release events occur more often in the S.R. region one week after KA-SE. The S.R. region is where labelled CA3 neurons would have their dendritic arbors and post-synaptic densities, which could suggest that UDP release occurs in more synapse-rich regions. While UDP1.0 was targeted to the neuronal membrane (hSyn promoter), this information merely discloses the site of detection, not the identity of release. Considering the importance of the “quad-partite synapse” formed between pre- and post-synaptic neurons, astrocyte fine processes, and motile microglial processes, potentially any cell could release UDP at this site. Future research is necessary to identify the mechanism(s) of release and their cellular origins. However, microglia are the only cells in the hippocampus and cortex that expresses the P2Y_6_ receptor necessary to sense UDP release^25,30^. Future development of a red-shifted UDP sensor will be integral for testing how well real-time UDP release *in vivo* can trigger microglial calcium signaling. As a first-generation sensor, UDP1.0 reports UDP release in a biologically relevant range (10 µM to at least 250 µM) and with high specificity, making it a valuable tool for future studies of UDP signaling dynamics, such as inflections^25^ and neurodegeneration^30-32^.

Our work indicates that cortical and hippocampal microglial utilize calcium signaling differently in response to regionally distinct environments. Microglial lysosome enrichment occurs in regions known to undergo cell death (amygdala, thalamus, and hippocampus). Transcriptomics, histology, and flow cytometry all indicate that P2Y_6_ activation (as an assumed response to UDP elevations) is a key driver of pro-phagocytic, pro-inflammatory transition in hippocampus. The fact that pro-inflammatory/pro-phagocytic transition does not occur in cortical microglia, despite the presence of P2Y_6_-based calcium signaling, suggests that calcium activity alone in microglia is not sufficient for reactive state transition.

Indeed, we were very surprised to see ameboid transition can also occur in microglia with strongly attenuated calcium signaling. We would hypothesize that neurons may provide microglia and other professional phagocytes with a coincidence of signals necessary for phagocytic transition and neuronal engulfment. Type I IFN activation may be a critical signal coinciding with P2Y_6_ pathway activation, as Type I IFN response genes can be enriched in *P2ry6^+/+^* microglia. Determining the timing and sources of Type I IFN in epileptogenesis and how it could engage in P2Y_6_ crosstalk is a key future direction. Beyond Type I IFN signaling, the P2Y_6_ pathway may engage in crosstalk with other damage recognition and phagocytic effector pathways in microglia. In particular, the Gas6-Axl signaling axis plays a key role in promoting phagocytosis in neuroinflammatory environments^53^. In general, while we have a decent understanding of which signals and contexts may activate different phagocytic effector pathways, we still lack a complete understanding of how the complex orchestration of phagocytosis occurs. Our work demonstrates that CD68 phagolysosome upregulation, necessary for processing synapses, toxic aggregates, pathogens, and even whole neurons, strongly depends on microglial calcium elevations. These finding may serve as critical context for why microglia engage calcium signaling during myelin sheath removal^54^, near Aβ plaques^55^, after seizures^5^, or in response to damage^4,7^. In future, it will be important to determine if other phagocytic receptor systems (*e.g.*, TREM2^56^, TYROBP, Axl, MerTK^53,57^, Spp1^58^) universally require calcium elevations for phagocytic function, particularly when lysosome enhancements may be necessary.

Finally, P2Y_6_-based immune signaling regulation may be critical in shaping the early epileptogenic or neurodegenerative landscape. Earlier studies have demonstrated that UDP or selective P2Y_6_ agonists can induce pro-inflammatory cytokine expression, but the cytokines created largely depend on the cell type and context. Activation of P2Y_6_ in blood lymphocyte culture preferentially stimulates IL-1β, TNFα, and IL-6^59^, while monocyte P2Y_6_ stimulation results in soluble TNFα and IL-8 expression^60,61^. Here we find that murine microglia in an epileptogenic environment increase their IL-1β and TNFα production when they have preserved P2Y_6_ signaling. Transcriptomic analyses suggest that P2Y_6_ signaling may ultimately arrive upon greater pro-inflammatory cytokine expression through NF-κB regulation. NF-κB represents a family of 5 different transcription factors typically sequestered in the cytosol^62^. Many of the *Trim* genes differentially expressed in P2Y_6_ KO mice regulate upstream elements of NF-κB activation, including poly-ubiquitination of TAB/TAK complexes or NEMO/IKK complexes necessary for NF-κB to enter the nucleus and begin cytokine transcription^37^. How P2Y_6_ signaling alternatively regulates *Trim* genes is an important future question of immune regulation, particularly given their inter-related functions in host-pathogen responses. On the other hand, calcium plays multiple known roles in IL-1β signaling, including its cleavage into an active form^63^, and secretion from myeloid cells^64,65^. We would therefore hypothesize that microglial P2Y_6_ calcium signaling could coordinate a pro-inflammatory phenotype at multiple levels, including Trim/NF-κB regulation of cytokine expression and calcium signaling for cytokine cleavage and release. Other studies suggest that P2Y_6_ phagocytosis and inflammation may not always be inter-related^66^. At the level of microglia, however, UMAPs suggest a strong overlap in the populations that perform phagocytosis (CD68^+^; NeuN^+^ inclusion) and express high levels of pro-inflammatory cytokines. Genotype-dependent differences in microglial IL-1β expression have perhaps the most promising implications for neuronal degeneration and seizure susceptibility. A wide literature^42^ has demonstrated that IL-1 antagonism can limit seizure severity^67^ or excitotoxicity^68^. Accordingly, exogenous IL-1 application can increase seizure severity^69^ or enhance NMDA calcium entry^70^, which may have important implications for cell death in the context of epileptogenesis^71^. The association between microglia performing phagocytosis and their high-level IL-1β/TNFα production may have important knock-on effects for neuronal health, particularly for neighboring neurons in dense pyramidal bands (such as CA3). We would posit that the known cytotoxic effects of these pro-inflammatory molecules may explain the continuing evolution of CA3 cell loss and prolonged FJC positivity observed in WT mice that is prevented by P2Y_6_ knockout. As such, P2Y_6_-associated cell loss may represent the combined impact of phagocytic clearance and additional immunopathic damage. Overall, our results suggest that microglial calcium activity can promote neurodegeneration through multiple coordinated mechanisms.

## Methods

### Animals

*P2ry6^-/-^* mice were kindly provided by Dr. Bernard Robaye of the University of Brussels (ULB). CalEx mice (R26^LSL-CAG-ATP2B2-mCherry^) were provided by Dr. Xinzhu Yu (University of Illinois Urbana-Champaign). C57BL/6J (WT) mice, Cx3Cr1^CreER-IRES-eYFP^ mice, TMEM119^2A-CreERT2^ mice, and R26^LSL-CAG-GCaMP6s^ mice were obtained from The Jackson Laboratory. Male and female mice were used in all studies, with procedures beginning at an age of 5 weeks. To mitigate the impact of background genetics in experiments, *P2ry6^-/-^* mice were crossed with in-house C57BL/6 mice to obtain *P2ry6^+/-^*. These mice were then intercrossed to generate *P2ry6^+/+^*and *P2ry6^-/-^* breeders for all non-imaging experiments. For microglial calcium imaging experiments, we similarly derived *P2ry6^+/+^*;Cx3Cr1^CreER-IRES-eYFP/WT^;R26^LSL-CAG-GCaMP6s/WT^ and *P2ry6^-/-^*;Cx3Cr1^CreER-IRES-eYFP/WT^;R26^LSL-CAG-GCaMP6s/WT^ lines from an intercross. Animals were group-housed on a 12h light-dark cycle in a temperature- and humidity-controlled room. To induce recombination, mice were weaned at P21 and given free access to tamoxifen in chow (500 mg tamoxifen/1 kg chow) for two- or four-weeks in their home cage. Tamoxifen-restricted littermates were separated at random during weaning and provided a standard diet during this period. Otherwise, assignment was based upon genotype and was therefore not randomized. However, investigators were blinded to genotype during experiments. Animal procedures were in agreement with NIH and Institutional Animal Care and Use Committee (IACUC) guidelines. Protocols were approved by the Mayo Clinic IACUC.

### Cell lines

HEK293T cells were obtained from ATCC. All cell lines were cultured at 37°C in DMEM (Biological Industries) with 5% CO_2_, supplemented with 10% (v/v) fetal bovine serum (FBS; GIBCO) and 1% penicillin-streptomycin (PS; Biological Industries).

### Molecular biology

To develop a UDP sensor, we inserted the cpEGFP module from GRAB_NE1m_ (containing flanking peptide linkers) into the chicken P2Y_6_ receptor. We screened TM5 insertion sites ranging from L214^5.65^ to V225^6.26^, and TM6 insertion sites from G221^6.23^ to M235^6.36^. All sensor variants were generated by overlap PCR without cloning into an expression plasmid. The forward primer is upstream of the CMV promoter-sensor-IRES-mCherry-CAAX-polyA expression cassette, amplifying a fragment containing the C-terminal half of cP2RY_6_ with IRES-mCherry-CAAX-polyA. The reverse primer targeted the TM5 insertion site using a 20-30 bp sequence homologous to the linker in the cpEGFP module from GRAB_NE1m_. This permitted the amplification of a fragment containing CMV promoter and N-terminal half of *cP2ry6*. A third DNA fragment was amplified containing the cpGFP module from GRAB_NE1m_. By performing overlap PCR, these 3 fragments were joined into a complete expression cassette. For the purpose of optimization, candidate sensor variants with known sequence alterations (*i.e.*, truncation variants, point-mutation variants) were sequentially screened. To optimize the signal (Figure S3A), we truncated the N-terminal linker of the cpEGFP module by 49 amino acids and the C-terminal linker by 2 amino acids. Additionally, four mutations were introduced at the interface between the cpEGFP module and the chicken P2Y_6_ sequence to produce GRAB_UDP1.0_. PCR reactions were conducted with a high-fidelity 2x PCR mix (Vazyme, P510-01). A final concentration of 1 M betaine and 1 mM DTT was added to the PCR mixture to improve specificity. Plasmids were generated using a Gibson assembly. DNA fragments were generated using PCR amplification with primers containing ∼25 bp overlap (XianghongShengwu, Beijing). All sequences were verified through Sanger sequencing by RuiboXingke (Beijing). The plasmids used to express the GRAB_UDP1.0_ sensor in HEK293T cells were based on the pDisplay vector, with an IgK secretion signal peptide added to the N-terminal of GRAB_UDP1.0_, following the IRES-mCherry-CAAX sequence, to ensure cell membrane labelling with RFP. The plasmid used to express the GRAB_UDP1.0_ sensor in mammalian neurons was cloned into a pAAV vector under the control of the hSyn promoter.

### Expression of GRAB_UDP_ variants in cultured cells

For the purpose of library screening, HEK293T cells expressing candidate GRAB_UDP_ sensor variants were seeded in 96-well plates (PerkinElmer). 1-2 μL PCR product mixed with 0.9 μL PEI MAX (Polyscience, 24765-100) was directly transfected into HEK293T cells in the 96-well plate. Following 12-hr transfection, the culture medium was replaced. Imaging or functional testing was performed after an additional 24-36 hours. For GRAB_UDP1.0_ *in vitro* characterization, 300 ng of plasmid was transfected per well into 96-well plates containing HEK293T cells following the time course of the library screening.

### Confocal imaging of cultured cells

Before imaging, culture media was replaced with Tyrode’s solution containing (in mM): 150 NaCl, 4 KCl, 2 MgCl2, 2 CaCl2, 10 HEPES, and 10 glucose (pH 7.3-7.4). HEK293T cells grown in 96-well plates were imaged using an Opera Phoenix high-content, confocal screening system (PerkinElmer) equipped with a 20x/0.4 NA objective, a 40x/0.6 NA objective, and a 40x/1.15 NA water-immersion objective, with 488-nm and 561-nm laser lines. Green fluorescence and red fluorescence were passed through 525/50-nm and 600/30-nm emission filters, respectively. Cells grown on 12-mm coverslips were imaged using a Ti-E A1 confocal microscope (Nikon) equipped with a 10x/0.45 NA objective, a 20x/0.75 NA objective, and a 40x/1.35 NA oil-immersion objectives, with 488-nm and 561-nm laser lines. Green fluorescence (GRAB_UDP_), and red fluorescence (mCherry-CAAX) were passed through 525/50-nm and 595/50-nm emission filters, respectively.

### Spectrum measurement

HEK293T cells were propagated in a 6-well dish, maintaining a seeding density of 2.6 x 10^5^ cells. Cells were then transfected using 4 μg of pDisplay-UDP1.0-IRES-mCherry plasmid. Following standard cell culture procedures, cells were detached using a 2.5% Trypsin solution and resuspended in 1 mL of DMEM with 10% FBS and 1% PS. Subsequently, cells were rinsed twice with Tyrode’s solution and resuspended in a final volume of 500 μL Tyrode’s solution. A 30 µL aliquot of cell suspension was placed into a 384-well microplate (Corning, CLS4514) for spectrum analysis using TECAN safire2. We recorded the excitation spectrum using a fixed emission filter at 540±10 nm, scanning the excitation wavelength from 385 nm to 520 nm with a 10 nm bandwidth filter. The emission spectrum was measured with a fixed excitation wavelength at 470 ± 10 nm, scanning the emitted light from 495 to 600 nm with a 10 nm bandwidth filter. To acquire the maximum fluorescence of GRAB_UDP1.0_, we added a final concentration of 100 μM UDP to the cell suspension. To acquire the minimum fluorescence, 5 U/mL of apyrase VII (Sigma) was added to deplete any potential basal UDP. Measurement of non-transfected HEK293T cells at a similar density served as a control. We recorded data from three wells for each condition.

### Luciferase complementation assay

The split-luciferase complementation assay was performed as previously described^72^. Briefly, 96-well cultured HEK293T cells were transfected with either cP2Y6-SmBit and LgBit-mGq (cP2Y6), UDP1.0-Smbit and LgBit-mGq (UDP1.0), or LgBit-mGq alone (Control). 24 hours after transfection, DMEM with 10% FBS and 1% PS was replaced by Tyrode’s solution. UDP (at concentrations ranging from 1 nM to 1 mM) was then applied with a final concentration of 5 μM furimazine (NanoLuc Luciferase Assay, Promega). Luminescence was subsequently measured using a Victor X5 multi-label plate reader (PerkinElmer).

### Induction of kainate status epilepticus (KA-SE)

Kainic acid was dissolved at 2 mg/mL in sterile saline (0.9%) and stored at room temperature for up to 3 weeks. Kainate was administered to mice without surgery at 5-9 weeks of age, or 3-5 weeks post-surgery. Mice received a first dose of 17.5 mg/kg KA (i.p.). Mice which did not display a Racine stage 4-5 seizure^73^ in the first hour were administered an additional 7.5 mg/kg (i.p.) injection every 30 min until a first Racine stage 4-5 seizure. After a first seizure, if mice did not display a Racine stage 4-5 seizure for 30 min, they were administered an additional 7.5 mg/kg injection. This process was followed until mice displayed 8 Racine stage 4-5 seizures. After the 8^th^ seizure, mice were administered valproic acid (250 mg/kg, i.p.) to attenuate SE. To mitigate the risk of batch variability and achieve more consistent neuropathology, we set inclusion/exclusion criteria based on initial seizure burden during SE. Specifically, mice needed to experience between 8-12 Racine stage 4/5 seizures in total. Mice above or below this range were excluded. Following valproic acid administration, behavioral seizure monitoring was discontinued once 30 min had elapsed from the timing of the last seizure.

### Viral Transfection

At least one hour prior to surgery, adult mice (5-8 weeks) were administered 5 mg/kg carprofen (s.c.) and 4 mg/kg dexamethasone (s.c.). Mice were then anesthetized with 3% isoflurane, shaved, and transferred into a stereotaxic frame (David Kopf Instruments), where an anesthetic plane was maintained with 1.0-2.0% isoflurane throughout surgery. To image longitudinal UDP release events *in vivo*, the skin overlaying the skull was cut away with scissors in a circular pattern and the pericranium was removed by peroxide. C57BL/6J WT mice were then injected through a drilled burr hole with AAV2/9-hSyn-UDP1.0 (titer: 2.9×10^12^ GC/mL), targeting layer 5 and layer 2/3 of somatosensory cortex (at 500 µm and 200 µm below the dura, respectively). A total of 250 nL AAV was dispensed at each cortical layer through a glass-capillary-attached Hamilton syringe at a rate of 40 nL/min using a UltraMicroPump-4 system (World Precision Instruments). To image UDP in hippocampus, we performed a midline incision of the scalp and performed stereotaxic AAV injection in C57BL/6J mice. We created a bilateral burr hole at AP: -1.8; ML:+/- 2.3 and slowly (1 mm/min) lowered a glass microcapillary to the level of CA3b (DV: -2.1 relative to Bregma). Both hippocampi were infused with 700 nL of AAV2/9-hSyn-UDP1.0 (1.5×10^12^ GC/mL) at 100 nL/min. After waiting 10 min post-infusion, the microcapillary was slowly raised and the midline incision was sealed with Vetbond (3M). To validate CalEx function, TMEM119^2A-CreERT2^; R26^LSL-CAG-ATP2B2-mCherry^ mice and TMEM119^2A-CreERT2^ mice were injected with rAAV-SFFV-DIO-GCaMP6f-WPRE-hGH-polyA (titer: 2.4×10^12^ GC/mL; Biohippo PT-6329). Bilateral injections were performed targeting the CA1 S.R. (500 nL at 100 nL/min; AP: -1.8; ML: +/-1.5; DV: -1.4 relative to Bregma). The midline incision was sealed with Vetbond. To ensure viral expression under the TMEM119^2A-CreERT2^ system, mice were fed tamoxifen chow for two weeks, beginning one week prior to viral transfection. In addition, a 100 mg/kg (i.p.) tamoxifen injection in corn oil was administered 3- and 5-days post-surgery.

### Cranial window surgery

For mice undergoing UDP1.0 transfection in cortex for *in vivo* imaging, the cranial window was installed during the transfection surgery. After UDP1.0 injection, a 3 mm circular craniotomy was created around the center of the injection site and the bone flap was carefully removed with fine forceps without tearing the dura. After bleeding was carefully resolved, a 3 mm circular coverslip (#0 thickness, Warner Instruments) was placed into the craniotomy and secured with blue-light curing dental glue (Tetric Evoflow, Ivoclar). Dental primer was then applied to the skull surface (iBond Total Etch, Kulzer) before a 4-point headpiece (Model 2, Neurotar) was secured in place with blue-light curing dental glue, sealing the entire skull surface. For studies of spontaneous calcium activity *in vivo*, CX_3_CR1^CreER-IRES-eYFP^;R26^LSL-CAG-GCaMP6s^ mice, which were either *P2ry6^+/+^* or *P2ry6^-/-^*, underwent cranial window surgery without viral transfection.

### *In vivo* 2-photon imaging

All *in vivo* imaging was performed at least 4 weeks after cranial window surgery (with or without viral transfection) to mitigate the impact of post-surgical inflammation and allow for robust viral transfection (if applicable). Prior to imaging, mice were acclimated to head fixation once daily for three days (15 min sessions) prior to the first imaging experiments. All imaging was performed in awake, head-fixed mice, which were able to ambulate on a floating platform (Neurotar Mobile Home Cage). Multiphoton imaging was conducted with a Scientifica Vista scope equipped with multi-Alkali photomultiplier tube detectors and galvo-galvo scanning. Excitation was achieved through a Mai Tai DeepSee laser, tuned to 940 nm for GCaMP6s or UDP1.0 imaging. Laser power under the objective was kept below 50mW during imaging. Green channel emission was passed through a 525 ± 50 filter. Spontaneous microglia calcium activity was captured as a 15 min T-series from a 300 x 300 µm area (512 x 512 pixels) at a 1Hz frame rate using a water immersion objective (Nikon, 16x; NA:0.8). A larger field of view (620 x 620 µm) and longer acquisition time (30 min) was used for UDP1.0 imaging. All *in vivo* imaging was performed 50-175 µm below the dura in somatosensory cortex.

### Brain slice preparation

Mice (6-9 weeks of age) were deeply anesthetized by isoflurane anesthesia (drop method, bell jar). After decapitation, the head was immediately cooled by submersion in ice-cold sucrose solution (185 mM sucrose, 2.5 mM KCl, 1.2 mM NaH_2_PO_4_, 25 mM NaHCO_3_, 25 mM glucose, 10 mM MgSO_4_, and 0.5 mM CaCl_2_, bubbled with 95% oxygen and 5% CO_2_ with an adjusted pH of 7.35-7.42 and an osmolarity of 280-290 mOsm/kg). The brain was then rapidly removed and 350 µm coronal sections were obtained via vibratome (Leica VT1000S), using the same ice-cold sucrose solution. Slices were rinsed in artificial cerebrospinal fluid (aCSF) before transfer to a recovery chamber containing 34°C aCSF (126 mM NaCl, 2.5 mM KCl, 1 mM NaH_2_PO_4_, 26 mM NaHCO_3_, 10.5 mM glucose, 1.3 mM MgSO_4_, and 2 mM CaCl_2_, bubbled with 95% oxygen and 5% CO_2_ with an adjusted pH of 7.35-7.42 and an osmolarity of 298-310 mOsm/kg). Slices were incubated at 34°C for 20 min then allowed to return to room temperature for 10 min prior to use. Slices were imaged using the same multiphoton system, equipped with a custom-made slice perfusion system (maintained at a 1-3 mL/min flow rate using room temperature aCSF). Reagents for extracellular solutions were acquired from Millipore-Sigma. All *ex vivo* imaging was performed at least 50 µm below the slice surface.

### Two-photon studies and analyses

#### Evaluation of spontaneous calcium activity

Spontaneous microglial calcium levels were assessed in *P2ry6^+/+^*;Cx3Cr1^CreER-IRES-eYFP^;R26^LSL-CAG-GCaMP6s^ and *P2ry6^-/-^*;Cx3Cr1^CreER-IRES-eYFP^;R26^LSL-CAG-GCaMP6s^ mice. Two T-series per mouse (900 fames) were captured from non-overlapping locations under the window and followed longitudinally for up to 14 days. T-series were processed in ImageJ by first correcting drift using subpixel registration through the Template Matching plugin. To establish somatic microglial ROIs, the user manually outlined these structures in an average intensity projection from the full field of view. Mean pixel intensity was captured from somatic ROIs using the multi-measure tool. Somatic ROIs were then masked in the average intensity image to decouple this region from microglial processes. Process ROIs could then be extracted from the average intensity image by thresholding the remaining image data and similarly using the multi-measure tool to capture mean pixel intensity in the T series. To mitigate selection bias, ROIs could only be selected from the average intensity image, which most strongly reflects the weak, constitutive eYFP fluorophore in microglia (CX_3_CR1^CreER-IRES-eYFP^) over GCaMP6s calcium dynamics. Mean pixel intensities for each ROI were converted to ΔF/F_0_ values through an Excel template by establishing the mean fluorescence for each of nine 100-frame intervals, then setting the F_0_ term as the lowest, non-negative 100-frame average. A calcium signal was considered significant at a ΔF/F level 3x greater than its standard deviation above baseline. The baseline is the average ΔF/F value in the lowest 100-frame interval while the standard deviation is calculated across all frames. Any and all frames above this threshold were summated to acquire the ROI’s calcium signal area (ΔF/F·s). For *ex vivo* spontaneous calcium imaging, the same genotypes and parameters (300 x 300 µm field of view, 1Hz, 15 min) were utilized. Spontaneous calcium activity in slice was processed and analyzed nearly identically to *in vivo* datasets, except for an additional correction step. Z drift which was minimal (microglial processes may shift but are not lost from the field of view; <4 µm) was corrected in the raw trace by fitting a linear slope to the drift and normalizing each frame against the slope term and frame number: F_corrected_ = F_frame_ - (frame · slope). This correction was not applied to ROIs which had a calcium event at the start or end of the recording. Z drift >4 µm, assessed immediately after acquisition, required video re-capture.

#### Evaluation of UDP release

After X/Y correction with the moco plugin, UDP events were manually identified from T-series by two independent investigators with strong congruency. UDP events were not uniformly radial, so their areas were outlined using the freeform tool to create ROIs, followed by ImageJ ROI area and X/Y-location (centroid) measurement. To obtain fluorescent values, mean fluorescent intensity was acquired through the multi-measure tool. To convert fluorescent intensity values into ΔF/F_0_ values, the F_0_ term was set as the median fluorescence in the 60 frames preceding event start. Event start was defined manually as the first frame preceding rise kinetics. The event end was calculated with a 10-frame moving window average. The first frame of the smoothed average which reached the F_0_ value was considered the event end. These frame values were used to define event time. A summation of ΔF/F values between these time points yielded the ΔF/F·s signal area for an event. A summation of ΔF/F·s signal areas from all events occurring within a 30 min acquisition was termed the cumulative signal area. *Ex vivo* UDP biosensor imaging occurred under identical settings (620 x 620 µm field of view, 1Hz, 15-30 min) for both the cortex and CA3 region. To induce epileptiform activity in slice, we perfused in daily aCSF lacking MgSO_4_. Both external solutions were corrected to have identical pH and an osmolarity within ±3 mOsm.

#### Evaluation of agonist calcium responses

Agonists were locally dispensed through a glass capillary at 3-5 psi for 300-600 ms using a Pico-spritzer III board (Parker Instrumentation). Agonists were kept in powder form (ATP and UDP) and dissolved/diluted at the time of experiment in daily aCSF with AF568 dye added for visualization (final concentration: 30 µM). Daily aCSF containing AF568 served as a negative control for pressure application. Red channel emission was passed through a 630 ± 75 filter (Chroma). To calculate an agonist response, a semi-automated script performed the following functions: (1) A threshold of AF568 dye spread in the red channel was used to limit the area of microglial ROI selection to cells likely encountering the agonist. (2) Within this area, an average intensity image was created of eYFP/GCaMP6s microglia from the first 30 frames (prior to agonist application) to select cells without bias towards response. A uniformly applied image threshold was used to automate microglia ROI selection using the analyze particles tool (usually representing a whole cell with soma and major processes). (3) Mean fluorescent intensities were gathered across 90 frames (30 baseline, 60 post-agonist application) for each ROI using the multi-measure tool. To calculate ΔF/F_0_ responses from the mean intensities, the F_0_ term was set as the mean of the first 30 frames (pre-agonist application). Significant calcium responses were considered as frames in the 60s post-agonist application period which were greater than the mean ΔF/F value of the first 30 frames plus 1.5x the standard deviation of the same period. Any and all frames above this threshold were summated across the 60 frames post-agonist application to acquire the signal area (ΔF/F·s). For dichotomous, “responding” and “non-responding” qualifications, an ROI needed to reach a signal area value above 10 ΔF/F·s.

#### Antagonist studies

Antagonist studies utilized a paired trial design in which UDP (500 µM) was first focally applied, then an antagonist was bath applied over a 10 min period before a second UDP focal application in the identical region. The antagonist effect was calculated as the overall average microglia ROI calcium response to UDP in the second (antagonist) trial divided by the first trial. Because repeated GPCR agonism can result in desensitization, antagonist effects were compared against the effect of desensitization, established by applying UDP twice in the same region, separated by 10 min over a series of independent trials.

#### Characterization of UDP1.0 ex vivo

To characterize UDP1.0 sensor affinity and selectivity, a pipette containing UDP or ATP was used to focally apply purines in the area of sensor expression at different concentrations (1 psi, 50 ms). In some trials, a longer duration of release was also performed (0.5 psi, 5 s). Florescent responses were calculated similarly to agonist calcium responses, except for a change in ROI creation. Eight square ROIs (7 x 7 µm), in a uniform configuration, were placed around the pipette tip, with the middle ROI centered on the pipette tip to acquire mean fluorescent intensities.

#### Validation of CalEx function

Evaluation of CalEx function was performed in naïve brain slices. In control mice (TMEM119^2A-CreERT2^ with tamoxifen treatment and rAAV-SFFV-DIO-GCaMP6f-WPRE-hGH-polyA transfection), 500 µM UDP was focally applied (3 psi, 200 ms) through a pipette with AF568 for visualization. Any microglial cell responding to UDP application in control tissue was analyzed using the same agonist-response analysis pipeline, except ROIs were manually created around responding cells. This was necessary because GCaMP6f transfection does not include an expression tag or produce a resting signal that is reliably detected above background levels. In CalEx mice (TMEM119^2A-CreERT2^;R26^LSL-CAG-ATP2B2-mCherry^), we similarly placed the UDP pipette in tissue; however, we acquired a 30 frame (1 Hz) image of the red channel at 1040 nm to clearly observe mCherry cell locations before UDP application/AF568 dye spread. We then applied 500 µM UDP from the agonist pipette in this local region. Similar to control tissue, we analyzed any cell responding to UDP application using a manual ROI for consistency. Because recombination could occur independently for CalEx-mCherry expression and GCaMP6f expression, only cells with a GCaMP6f calcium response and an mCherry signal were included for analysis in the CalEx group.

#### Chemotaxis studies

Chemotaxis studies were performed near L2/3 of cortex by inserting a 1 mM ATP- or UDP-containing pipette 45-60 µm into the slice with an AF568 dye in the pipette for visualization (50 µM). Process movement in *P2ry6^+/+^*;CX_3_CR1^CreER-IRES-eYFP^;R26^LSL-CAG-GCaMP6s^ and *P2ry6_-/-_*_;_CX_3_CR1^CreER-IRES-eYFP^;R26^LSL-CAG-GCaMP6s^ slice was visualized via the weak eYFP tag over a 20 min trial (1 Hz imaging, 150 x 150 µm area). For analysis, a 60 s frame average of the eYFP signal was gathered for each of the 20min periods. The location of the pipette tip was recorded as a fixed point in X/Y-space. Processes moving towards the pipette tip (starting from within 40 µm) were tracked per video by their bulbous tip location (in X/Y space) across the 20 average frames. The distance from pipette was calculated as the hypotenuse length formed in X/Y space from process tip to pipette tip (a=X_pipette_-X_process_; b=Y_pipette_-Y_process_; distance=√a^2^+b^2^). All agonists and antagonists were acquired from Tocris. Experiments were discontinued 3.5 h after slicing.

### Tissue Collection and Immunofluorescence

After terminal isoflurane exposure (drop method, bell jar), mice were transcardially perfused with ice-cold 1x PBS for exsanguination (30 mL) followed by 50 mL of ice-cold 10% neutral-buffered formalin (Labchem LC146706). Whole brains were post-fixed overnight at 4°C in 10% neutral-buffered formalin, followed by 30% sucrose in PBS for cryoprotection. Coronal sections (20 µm) were cut by a cryostat (Leica CM1520) to capture sections from dorsal to ventral hippocampus. Sections were stored at 4°C in 1x PBS containing 0.01% sodium azide. For immunofluorescent staining, sections were plated onto slides, then blocked and permeabilized for 1h using 0.4% Triton X-100 in CytoQ immuno-diluent (Vector Biosciences). Sections were then incubated overnight at 4°C with primary antibodies in CytoQ containing 0.4% Triton X-100: anti-IBA1 (1:500 Rb for Sholl analysis; 1:400 Gt for phagocytosis), anti-NeuN (1:500 Rb for neuron survival; 1:500 Ms for phagocytosis), anti-CD68 (1:500 Rb), anti-IL-1β (1:400 Ms), and anti-cFOS (1:500 Rb). After primary antibody incubation, sections were washed with 1x PBS. Sections were then incubated at room temperature for 1.5 hours with secondary antibodies in PBS containing 0.4% Triton X-100 (1:1000, Goat or Donkey; Life Technology). Secondary antibodies were washed with 1x PBS, counterstained with DAPI (1 µg/mL, 10 min), and mounted with Fluoromount-G (Thermo).

### FluoroJade-C (FJC) Staining and Analysis

For FJC staining, sections were plated on to slides for batch processing in 50 mL Coplin jars. Sections were first exposed to 80% ethanol with 5% NaOH, followed by 70% ethanol, then ultrapure water. Sections were blocked for 5 min with 0.04% KMnO_4_ in water, then exposed for 20 min to FJC (0.0001% FJC working solution diluted in 1% acetic acid with 1 µg/mL DAPI). Sections were then washed and mounted with DEPEX (Electron Microscopy Sciences) for imaging. FJC staining was imaged using an EVOS FL fluorescent microscope (Thermo Fisher Scientific). Bilateral images of the CA3 pyramidal band were taken at 3 different levels (sections) of dorsal hippocampus per animal and averaged per hemisphere (10x objective, 1024 x 768 pixels). Average FJC area values were established by uniform threshold and restricted to a 400 x 400 µm area centered on the pyramidal band.

### Sholl Analysis

Microglial Sholl analysis was performed using fixed-tissue imaging of IBA1 staining in dorsal hippocampal CA3 and somatosensory cortex. Imaging was performed with a Zeiss LSM 780 confocal microscope equipped with a 40x water-immersion objective (NA: 1.2). The DAPI channel was used to visually center the imaging field on the CA3 region or layer 2/3 of cortex, near the midline. A Z-stack volume was then acquired at 512 x 512-pixel resolution (212 x 212 µm) across a uniform 14 µm stack with a 1 µm step size. An average intensity image of the IBA1 channel across Z levels was then used for Sholl analysis. Sholl morphology was calculated using an ImageJ plugin^74^, with the center of the soma being user defined. Thereafter, intersections were calculated along the path of concentric circles starting 6 µm from the soma center and increasing by 1 µm in radius. Longest process length was defined by the largest concentric circle capturing a process ramification. Total intersection values were defined as the sum of all process intersections from Sholl radii for a cell.

### CD68 imaging and analyses

To determine CD68 area across brain regions, sections were imaged with a Keyence BZ-X810 fluorescent microscope in the amygdala, dorsal and ventral thalamus, and hippocampus (CA3 and CA1). Images were taken from a single Z-plane (1449 x 1091 µm) with a 10x objective. A uniform threshold was applied to each image in ImageJ to determine CD68^+^ area. CD68 expression levels in CalEx tissue were analyzed from average-intensity LSM 780 confocal Z-stack images (1024 x 1024-pixel resolution with 1 µm Z steps; 20x, NA:0.8). To determine CD68 expression area in CalEx-mCherry positive and negative microglia, a code performed the following steps: (1) A uniform threshold of the neutral IBA1 channel was created and established ROI outlines around IBA1 cells. (2) A uniform threshold of the CD68 channel was created. (3) Each IBA1 ROI outline is overlaid on top of the CD68 threshold image. The CD68 area captured within each IBA1 outline is recorded as the raw area and the normalized to the IBA1 outline (% area). (4) To determine if these CD68 values occur in an mCherry^+^ or mCherry^-^ cells, a uniform threshold of the mCherry image was taken and each IBA1 ROI was then overlaid on top of the mCherry threshold image. Any IBA1 outline which has 67% area overlap with mCherry is considered CalEx positive. Because the mCherry tag is cell-filling and can illuminate smaller microglial processes, 35-40% of microglia are counted as mCherry positive, matching recombination rates. mCherry^-^ microglia are also clearly distinct, with 50-60% of microglia in an imaging plane having 10% or less area overlap between mCherry and IBA1. Typically, under 15% of cells fall into an intermediate range of IBA1/mCherry overlap, suggesting these criteria can effectively separate the two populations without user bias. To determine CD68 overlap with NeuN neurons in the CA3 region, a similar analysis pipeline was used. (1) A uniform threshold of the NeuN channel was created within the pyramidal band to establish individual NeuN ROI outlines. (2) A uniform threshold of the CD68 channel was created. (3) Each NeuN ROI outline was sequentially overlaid on top of the CD68 threshold image. The CD68 area captured within each NeuN outline was recorded as the raw area and the normalized (%) area of the NeuN outline. (4) The percent overlap was then segmented into no/minimal overlap (<10% area overlay), moderate interaction (between 10-80% overlap), and engulfment (>80% overlap). The same pipeline was applied to the IBA1 channel in place of the CD68 channel for additional analysis.

### NeuN analyses

To determine neuronal cell density, NeuN sections were imaged with a Keyence BZ-X810 fluorescent microscope in the CA1 and CA3 regions of hippocampus. A single Z-plane image was taken with a 20x objective. A uniform threshold was applied to the NeuN channel and the watershed tool was used to separate overlapping nuclei. A 100 x 100 µm ROI was evenly placed over 6 different locations of the pyramidal band per image. The number of NeuN somata counted by ImageJ (analyze particles) within this ROI was averaged across the 6 sampling areas to obtain a single value per mouse. An identical approach was used to quantify Cresyl Violet staining. To determine NeuN intensity profiles in CA3, three parallel lines were drawn along the CA3 pyramidal layer (one nearer to S.R., one in the middle, and one nearer to S.O.). An intensity profile along these lines was computed using the multi-plot tool and averaged to obtain a single profile per section. These profiles are only intended for illustrative purposes in representing regions of cell loss and were not formally quantified.

### Imaris rendering

Three-dimensional renderings of DAPI/NeuN/CD68/IBA1 interactions were performed by imaging a representative Z stack with a 63x objective (oil, NA:1.4) and a 0.33µm step size on a Zeiss LSM 980 confocal microscope. The airy scan module was used for idealized fluorophore detection at 1884 x 1884- pixel resolution (66.5 x 66.5 µm). Rendering was performed in Imaris. IBA1 microglia and NeuN neurons were manually surface rendered, while CD68 lysosomes were represented as center-aligned spots of growing size (>0.5 µm) based upon absolute intensity.

### High-parameter flow cytometry and analysis

Tissue was harvested in two cohorts, following a pilot experiment to determine panel viability and establish ideal antibody titers. Post-SE tissue was harvested 5 days after KA-SE, while a naïve group was established by enrolling saline-injected sibling littermates from both genotypes in each experimental batch. After terminal isoflurane exposure, mice were exsanguinated using ice-cold 1x PBS solution to remove cell populations transiting the vasculature. The brain was rapidly removed and mechanically dissected to obtain the full cortex and bilateral hippocampi. Remaining brain tissue was utilized for staining controls. Tissue was enzymatically digested in C tubes for 30 min using the tumor dissociation kit with mild mechanical movement and heating at 37°C (Miltenyi Octo Dissociator: adult brain dissociation protocol). Tissue was further processed into a single-cell suspension following manufacturer’s instructions, including steps to remove debris and myelin. Samples were then exposed to a viability dye (1:1000 Zombie UV) for 20 min at RT. Samples were washed in PBS and exposed to a surface antibody panel for 30 min at 4°C, including CD45-PerCP (1:50), CD11b-PE/Cy5 (1:100), CD126-BUV563 (1:100), MERTK-BV786 (1:100), CD25-BV750 (1:100), CX3CR1-PacificBlue (1:100), P2Y12-APC-Fire810 (1:100), TCRβ-PE/Cy7 (1:100), CD11c-SPARK-NIR-685 (1:100), Ly6G-BV711 (1:200), CD4-BV510 (1:50), CD8α-BV570 (1:50), CD122-BB700 (1:100), F4/80-BUV395 (1:200), NK1.1-BUV615 (1:100), MHCII-BUV805 (1:200). After additional wash steps, samples were fixed and permeabilized by 20 min exposure to Cytofix/Cytoperm solution (BD #554722) at 4°C. Samples were then incubated with an intracellular antibody panel for 30 min at RT in 1x permeability/wash buffer (BD #554723), including NeuN-APC (1:50), Vimentin-CL488 (1:50), TNFα-BV650 (1:100), Pro-IL1β-PE (1:50), CD68-BV421 (1:50), STING-PE/CF594 (1:50), IL2-BV605 (1:50), IFNγ-BUV737 (1:50). After a final series of wash steps, samples were kept at 4°C and analyzed within 24 hours using a 5-laser Cytek Aurora spectral cytometer. For an antigen, the level of staining considered as true biological signal was established with a fluorescence-minus-one (FMO) control in which all fluorophores in the panel except for the fluorophore of interest were used in staining. Gating strategies are described in Figure S8. Due to batch-associated variability in antibody signals between cohorts, frequency calculations are normalized against the average naïve value per cohort. Analyses were performed in FloJo.

### Transcriptomic Analysis

For bulk RNA-seq, microglia were isolated from the brains of *P2ry6^+/+^*and *P2ry6^-/-^* mice three days after KA-SE. Briefly, mice were transcardially perfused with ice-cold PBS, and the olfactory bulbs and cerebellum were removed from the brain. Remaining brain tissue was dissected into small pieces (∼1 mm^3^) in ice-cold PBS. Tissue was then processed by enzymatic digestion following the manufacturer’s instructions (Miltenyi Adult Brain Dissociation Kit), including debris removal. Red blood cell removal was omitted due to perfusion. Single-cell suspensions were labeled with CD45-PerCP (1:100) and CD11b-BV650 (1:100) and sorted by a BD FACS Aria to obtain CD45^Mid-Lo^;CD11b^Hi^ microglia, immediately resuspended in Trizol. Isolated RNA quality was assessed with an Agilent 2100. Qualifying samples (N=3 *P2ry6^+/+^* and N=4 *P2ry6^-/-^*) with an RNA integrity number (RIN) >7 were processed for library preparation with the SMART SEQ II kit in a single batch. Pair-ended sequencing (PE, 100bp) was performed with the BGISEQ platform. Cutadapt was used to remove adapters from the paired-end raw reads^78^. Trimmed, paired-end reads were aligned to the mus musculus reference genome (mm10) using STAR^75^ (v.2.5.4b). Evaluation of DEGs utilized the DESeq2 (v1.30.1) pipeline^76^ with raw read counts estimated from STAR. The gene set provided as an initial input contained approximately 13,700 genes with a base mean value >0.5 to filter out virtually undetected transcripts. No other filtering was performed and default settings were used in the DESeq2 pipeline. Evaluation is presented as a WT – KO contrast, where transcripts higher in the WT mouse have a positive log_2_FC value. DEGs include genes with a log_2_FC>±1.0 and a P_adj._>0.1. Of note, RSEM^77^ (v1.3.1) was used to calculate transcripts per million (TPM) from the .bam files generated by STAR.

### Novel Object Recognition

Novel object recognition (NOR) tasks were performed in a cohort of *P2ry6^+/+^* and *P2ry6^-/-^* mice in a naïve baseline state and one week after KA-SE. The arena consisted of a (42 x 42 x 25 cm) plexiglass box with a consistent symbol placed on each wall as a spatial cue. The entire plexiglass arena was enclosed within an insulated box providing temperature and humidity control as well as consistent lighting. Behavior was monitored by an overhead camera system. Mice were habituated to the arena for 5 min/day for 3 days preceding the baseline test. Habituation was also performed one day before the KA-SE test. The test consisted of a first “object familiarization” phase where two identical objects were placed along the center axis of the arena spaced 6.5 cm from the left (“A”) and right (“B”) wall. Mice were allowed to explore the arena and objects for 5 min. Mice were returned to their home cage for a 30 min period before a “novel object” phase began. In the novel object phase, a familiar object was placed in a previous location (A or B) and a novel object was placed in the other location. The identity of the novel object (between two different object sets) and the location of the novel object (A/B) was randomized across trials. Mice were given 5 min to explore the novel and familiar object in the novel object phase. The objects and arena were thoroughly cleaned with 70% ethanol after every 5 min trial. In a second NOR test performed 7 days after KA-SE (10 days after the first test), the same procedures were followed. Notably, two completely different object sets were used in the day 7 post-SE test. During the object familiarization phase, a preference index was established as the time spent exploring an object in location A divided by the time spent exploring the identical object in location B. In the novel object phase, a discrimination index was determined as the time spent exploring the novel object divided by the time spent exploring the familiar object. Exploration time was established by a consistent reviewer blinded to genotype. Exploration time was counted when mice were within 3 cm of the object and either facing the object smelling the object, or contacting the object. Bouts in which mice attempted to climb or scale the object were not counted as object exploration. Additional analysis was performed using the open-source EZtrack program to track mouse location, report overall mouse motility in a trial, and determine periods in which mice were within the novel or familiar object vicinity (by a consistent, user-defined ROI).

### Statistics and Quantification

All statistical analyses were performed in GraphPad Prism. Before running statistical testing, the normality and variance of the distribution was determined to arrive on the appropriate characteristics for a statistical test. A t-test was used to compare two groups at a single timepoint. To compare two groups across time, a Two-way ANOVA design was used with either a Sidak’s post-hoc test (comparison between genotype/group at each time point), or a Dunnett’s post-hoc test (comparison of a time point against the within-group baseline). A repeated-measures design was only used for *in vivo*, longitudinal datasets comparing the same region over time. Comparing ≥3 groups/categories at a single time point utilized a One-way ANOVA with either a Fisher’s post-hoc test (comparison to baseline), or Tukey’s post-hoc test (comparisons between all groups). All post-hoc testing accounted for multiple comparisons. Correlations between seizure severity and outcome measures were reasonably well fit by a simple linear regression, with additional slope and intercept comparisons. Neuropathology outcomes from mice not reaching seizure inclusion criteria were processed for the sole purpose of correlational studies to achieve a wider range of outcomes. Responses to UDP across a concentration range were fit by a nonlinear log(agonist) vs. 3-parameter response equation in Prism. Sample sizes were based on prior experience using similar techniques^5^. Reduced neuropathology outcomes in *P2ry6^-/-^* mice were confirmed across two cohorts at the day 1, 3, and 7 time points, with similar findings. Experimenters were blinded to genotype for *in vivo* imaging studies, histology studies, and behavioral studies. Analysis was either performed with blinding to genotype (behavior) or using automated analysis pipelines which strongly mitigate user bias.

## Author Contributions

A.D.U and L-J.W. conceptualized the study; A.D.U., B.L., K.A., Y.Li, and L-J.W. designed methodology. B.L., Z.W., and Y.Li designed, optimized, and validated the UDP sensor. A.D.U., S.Z., Y. Liang, and M.X. conducted *in vivo* experiments. K.A. conducted flow cytometry studies and analyses. A.D.U. and G.T. performed histology and image analysis. J.Z. and S.Z. performed isolation procedures for Bulk-seq. B.H. and J.S. analyzed transcriptomic data. A.D.U. and L-J.W. wrote the manuscript within input from all authors. X.Y., A.J.J., Y.Li and L-J.W. provided reagents and equipment.

## Acknowledgements

We thank Dr. Bernard Robaye (Universite Libre de Bruxelles) for the *P2Y6^-/-^* line. We also thank Alexander Anwar and the Mayo Microscopy and Cell Analysis Core (MCAC) for assistance with image rendering. Institutional access to Imaris and BioRender was kindly provided by the Mayo Foundation. This work was supported by grants from the NIH (K99 NS126417 to ADU; R01 NS103212 and RF1 NS122174 to AJJ; R01 NS088627 and R35 NS132326 to L-J.W.) and the National Natural Science Foundation of China (31925017 to Y.Li). The funders had no role in study design, data collection and analysis, decision to publish or preparation of the manuscript.

## Declaration of Interests

The authors declare no competing interests.

## Supplemental Figures

**Figure S1:**
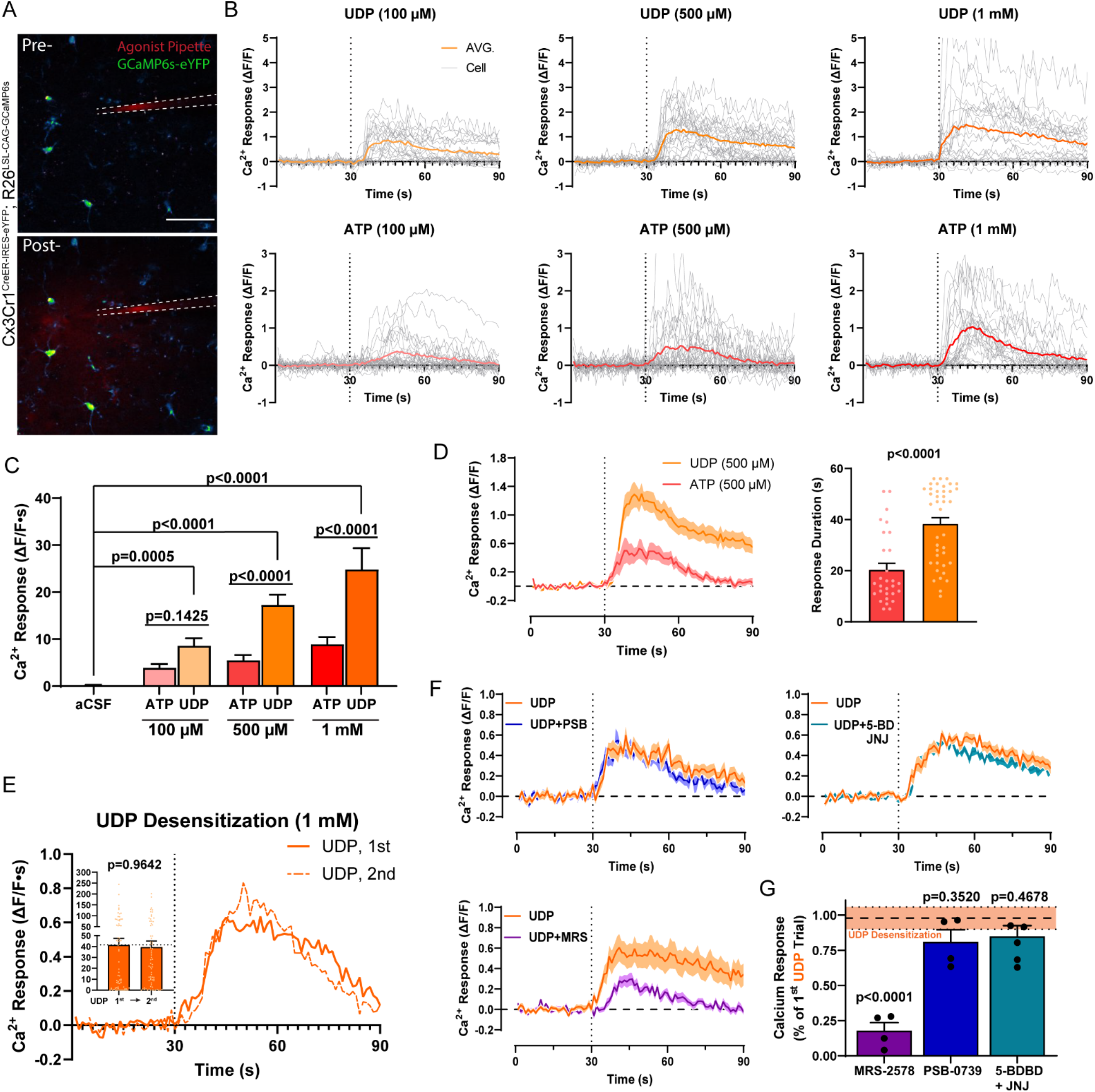
UDP signals through P2Y_6_ receptors to evoke calcium signaling. (A) Image of an acute brain slice prepared from mice expressing GCaMP6s in microglia. An agonist is focally applied through a glass capillary with AlexaFluor 568 dye for visualization. Scale bar, 100 µm. (B) Representative microglial calcium responses (ΔF/F) to ATP or UDP application (dotted line represents time of agonist application). (C) Overall calcium responses in naïve microglia to aCSF control or purine application. Two-Way ANOVA with Dunnett’s post-hoc comparison between aCSF and UDP, or Sidak’s post-hoc test for UDP vs ATP (4-6 slices prepared from N=3-4 mice;). (D) 500 µM UDP or ATP calcium response duration (from paired trials in 3 slices from N=3 mice; dots represent ROIs; unpaired T-test). (E) Calcium responses to repeated UDP application to determine the effect of desensitization (line graph: average ΔF/F calcium responses to UDP from a representative paired trial; bar graph: overall UDP calcium response across paired trails in 4 slices from N=4 mice; dots represent ROIs; paired T-test). (F) Calcium responses to UDP alone and in a paired trial with bath-application of a selective antagonist (MRS-2578 (MRS, 40 µM): P2Y_6_ antagonist; PSB-0739 (PSB, 5 µM): P2Y_12_ antagonist; 5-BDBD (5-BD, 20 µM): P2X_4_ antagonist; JNJ-54175446 (JNJ, 10 µM): P2X_7_ antagonist). (G) Summary of antagonist effects on UDP calcium responses. One-Way ANOVA with Dunnett’s post-hoc comparison to desensitization (orange horizontal line represents desensitization mean ± SEM; dots represent an individual trial in one slice per mouse). All bar graphs or shaded line graphs represent the mean ± SEM.

**Figure S2:**
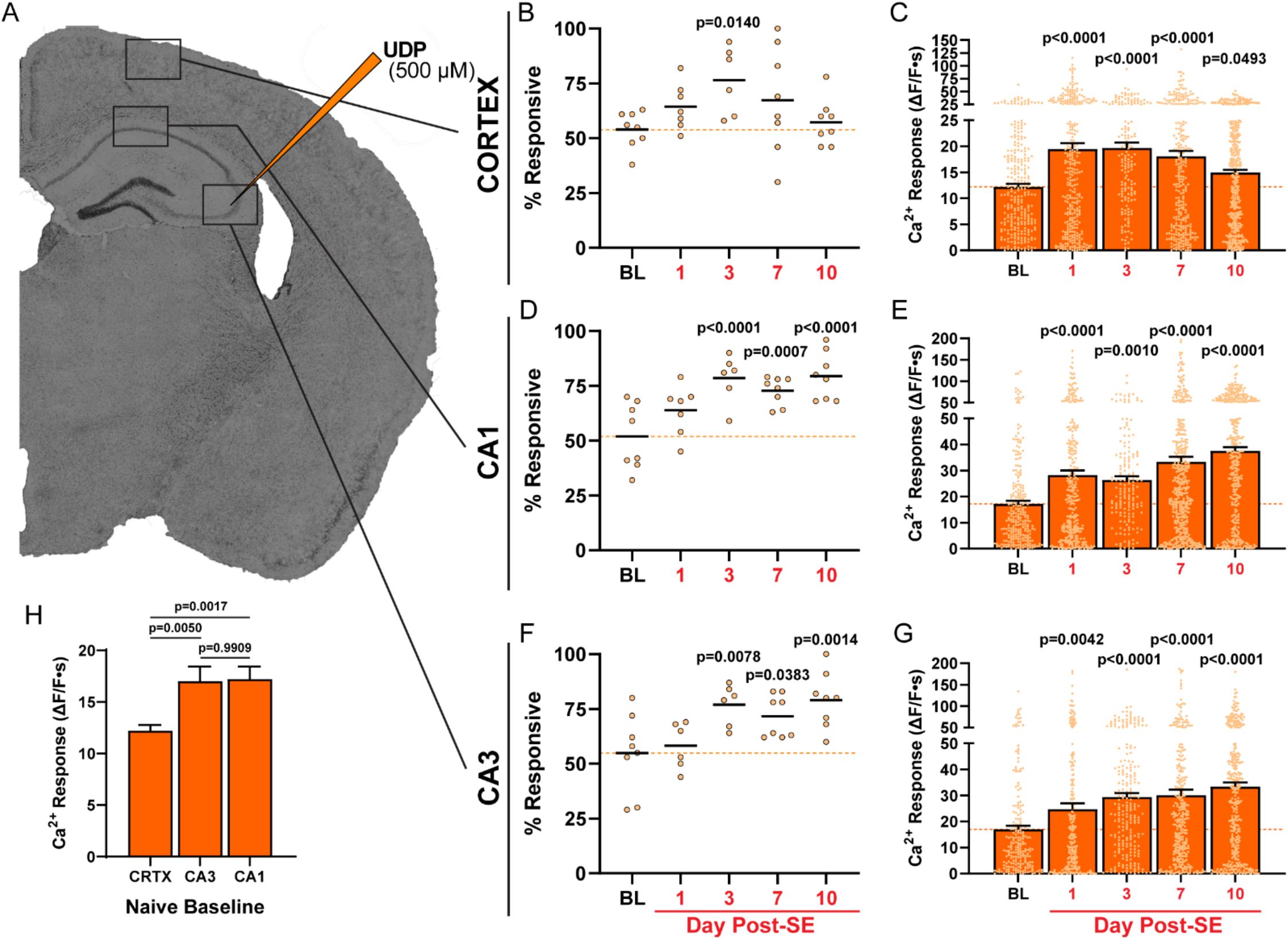
UDP calcium responses and spontaneous activity increase during early epileptogenesis in slice. (A) Regions surveyed in acute brain slice for microglial calcium responses. (B, D, F) The percentage of microglial ROIs responding to 500 µM UDP application in cortex (B), CA1 (D), or CA3 (F). Dots represent an individual slice (2 per mouse, N=3-4). Line is at the mean. One-Way ANOVA with Dunnett’s post-hoc comparison to baseline. (C, E, G) The calcium response from microglial ROIs after 500 µM UDP application in cortex (C), CA1 (E), or CA3 (G). Dot represents an individual ROI (from 2 slices per mouse, N=3-4). One-Way ANOVA with Dunnett’s post-hoc comparison to baseline. (H) Differences in baseline (naïve) microglial calcium responses to UDP between brain regions. One-Way ANOVA with Tukey’s post-hoc comparison (data from all ROIs surveyed across 2 slices per mouse, N=4). Bars represent the mean ± SEM.

**Figure S3:**
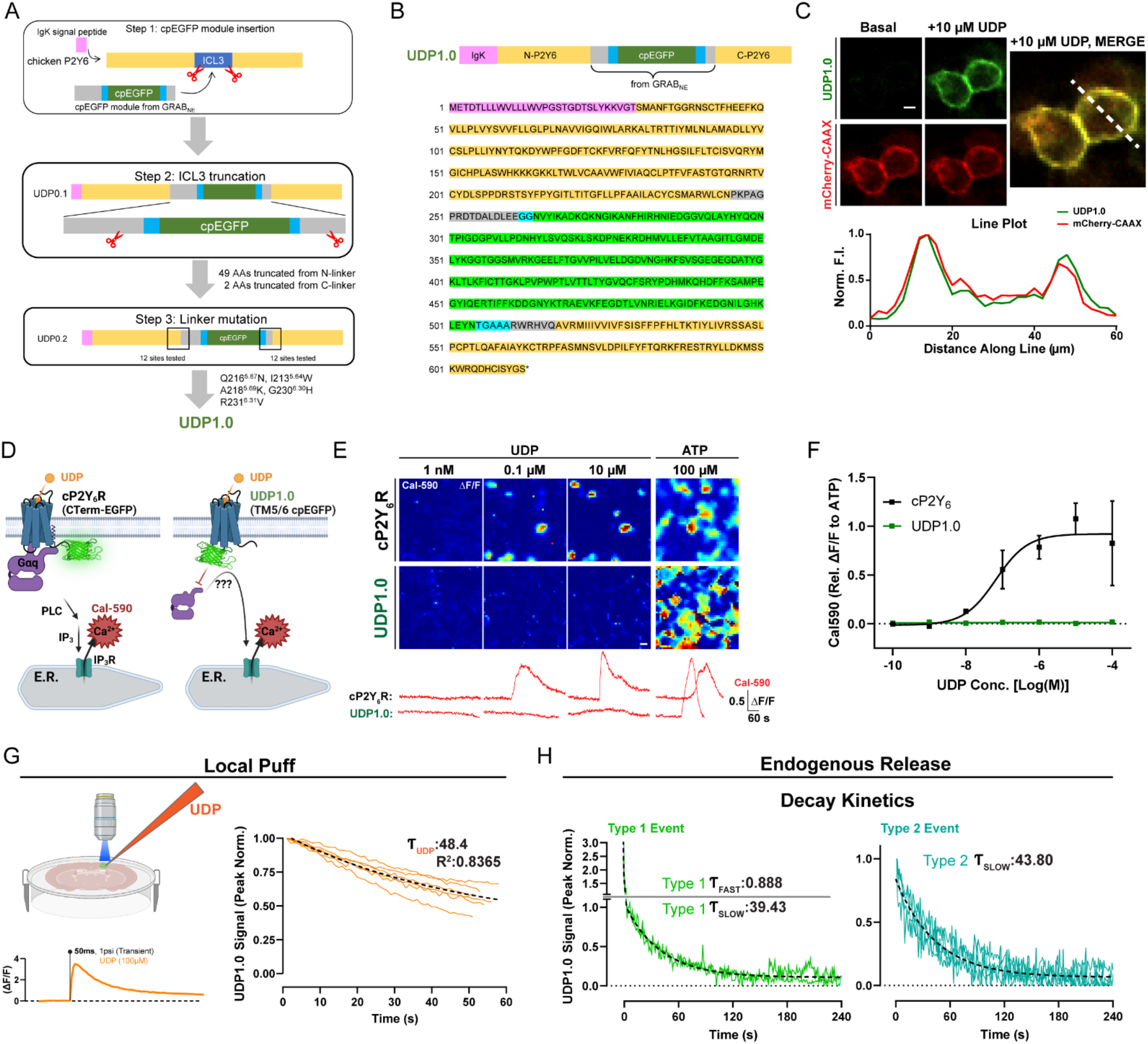
UDP sensor sequence, coupling, and kinetics, related to Figures 1 and 2. (A) Steps taken to optimize and refine UDP sensor signal. (B) Amino-acid sequence of UDP1.0. (C) Evaluation of UDP1.0 sensor trafficking to the membrane with membrane-trafficked mCherry-CAAX expression for reference. (D) Experimental schematic: Calbryte-590 calcium imaging in response to purines using HEK cells transfected with cP2Y_6_ or UDP1.0. (E) Cal-590 calcium response images and corresponding ΔF/F traces from cP2Y_6_- or UDP1.0-transfected HEK293T cells in response to purines. (F) ATP-normalized Cal-590 fluorescent responses across a range of UDP concentrations (mean ± SEM; lines represent a non-linear fit of log (agonist) vs. 3-parameter response). (G) Schematic of *ex vivo*, local UDP application in a UDP1.0-transfected brain slice with a representative ΔF/F response. Peak-normalized responses were used to determine the UDP1.0 single-component ͳ decay value in response to UDP (12 trials from 3 slices and mice). (H) One- or two-component ͳ decay fits for Type 1 and Type 2 UDP events observed *in vivo* (ͳ value reflects n=15 Type 1 and n=22 Type 2 events aggregated across N=5 mice).

**Figure S4:**
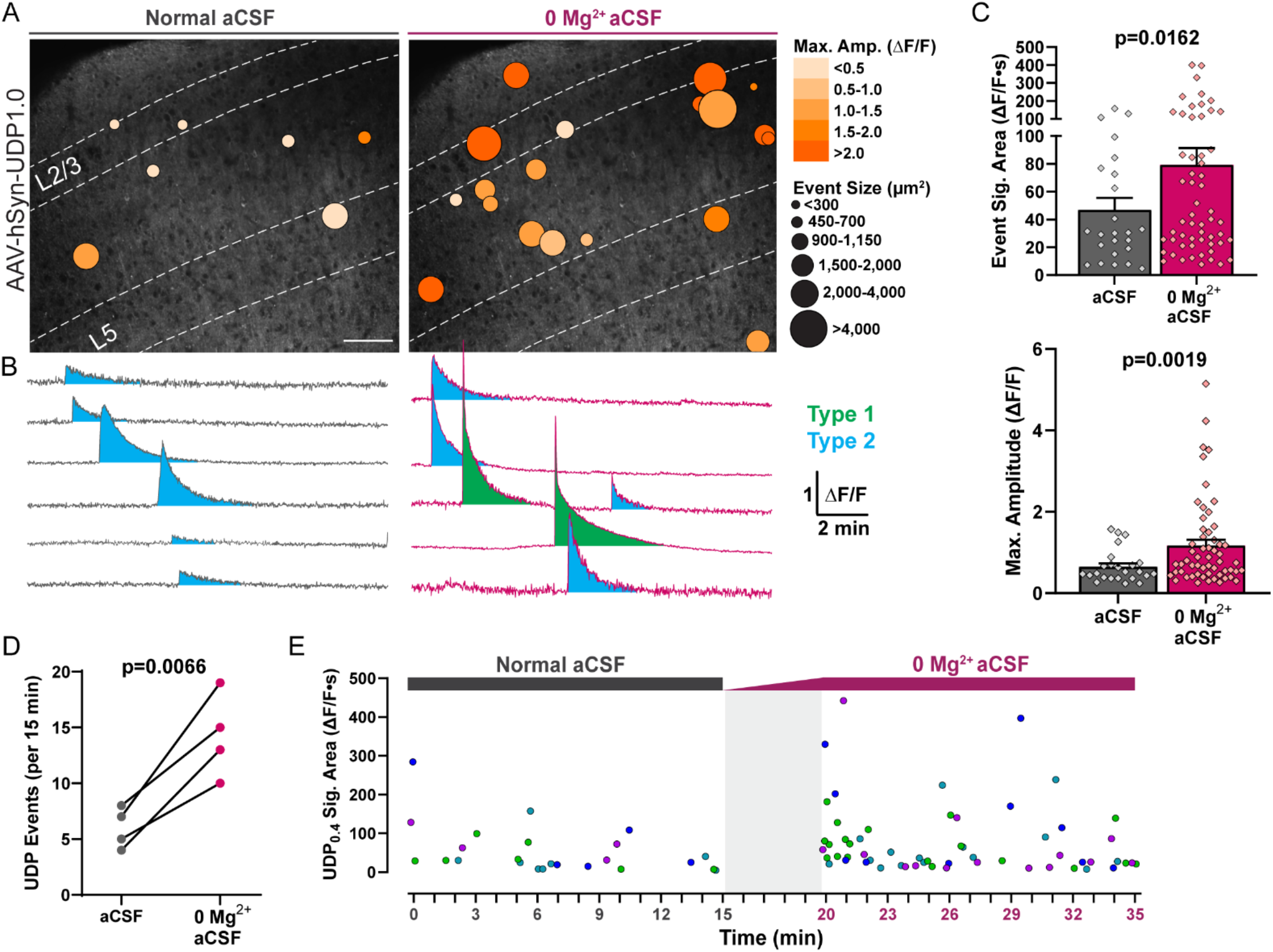
Enhanced UDP release during 0 Mg^2+^ hyperexcitability, related to Figure 2. (A) UDP1.0 sensor events overlaid across cortical layers during bath incubation with normal aCSF or 0 Mg^2+^ aCSF. Scale bar, 100 µm. (B) Event traces (ΔF/F) demonstrating Type 2 events in the presence of normal aCSF, or a combination of Type 1 “sharp peak” events and Type 2 events during 0 Mg^2+^ conditions. (C) Comparison of UDP event signal area and maximum amplitude between normal and 0 Mg^2+^ aCSF conditions. Student’s t-test (dot: one event; bar: mean ± SEM). (D) Paired UDP event frequency under normal and 0 Mg^2+^ aCSF conditions. Paired ratio T-test (one slice from N=4 mice). (E) Plot of UDP event signal area (ΔF/F·s) and timing over 15 min of normal aCSF incubation or 15 min of 0 Mg^2+^ aCSF incubation (gray box: 5 min solution exchange period; dot: one UDP event where color corresponds to a trial; one slice from N=4 mice).

**Figure S5:**
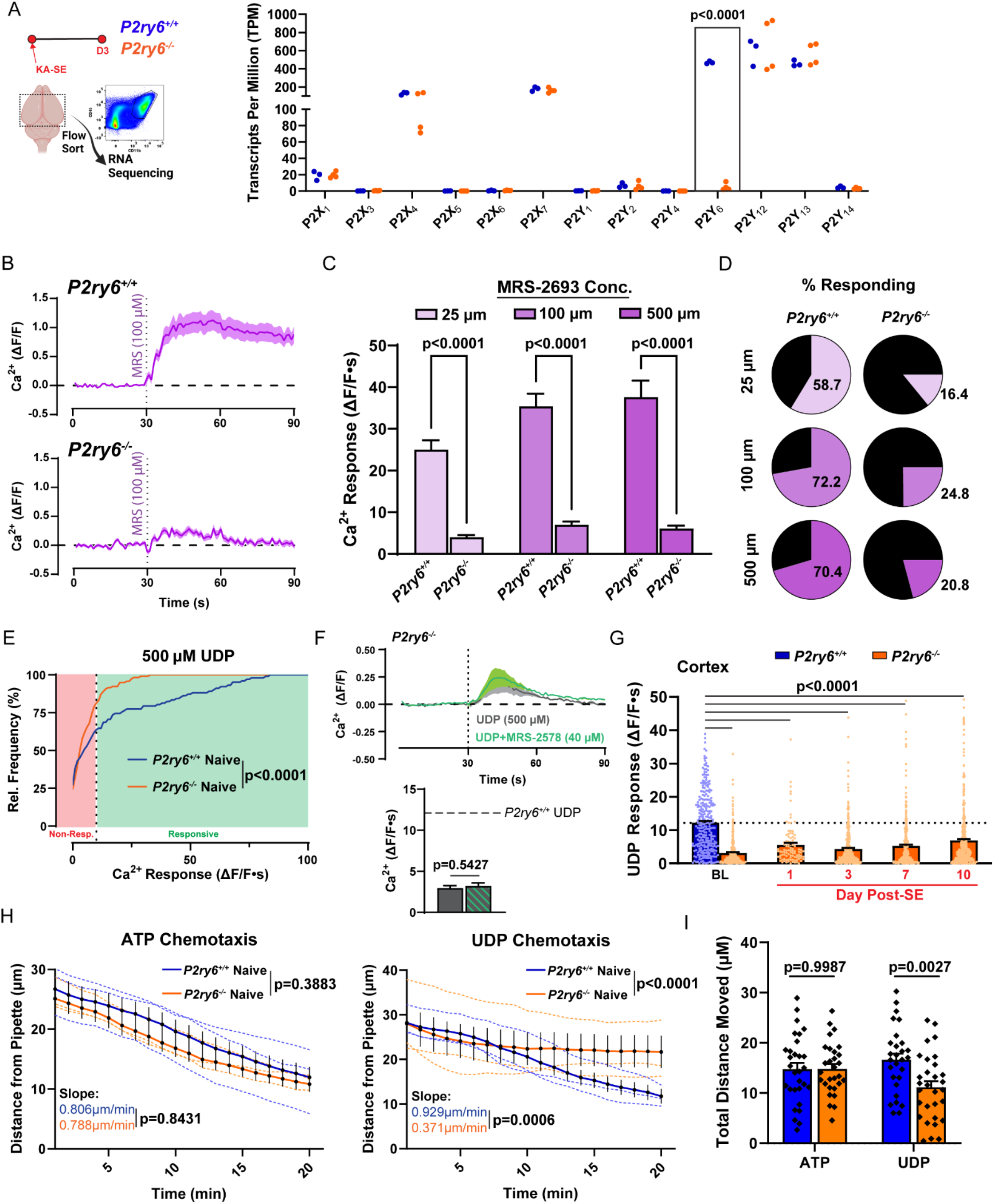
Functional characterization of the *P2ry6^-/-^* mouse, related to Figure 3. (A) Isolation of microglia for transcriptomic analyses 3 days after KA-SE. Confirmation of P2ry6 transcript loss from brain microglia without changes in other P2 receptors transcripts. Two-Way ANOVA with Sidak’s post-hoc test (dot: one mouse/sample). (B) Representative *P2ry6^+/+^* and *P2ry6^-/-^* microglial calcium responses to MRS-2693, a high affinity P2Y_6_ agonist, in naïve brain slice. (C) Overall calcium responses to MRS-2693 across a concentration range. Two-Way ANOVA with Sidak’s post-hoc test (all concentrations tested in 2-3 trials per slice; 4 slices from N=4 mice per genotype). (D) Percentage of microglia responding to MRS-2693. (Average across trials; same dataset as C). (E) Distribution of microglial calcium responses to 500 µM UDP by genotype. Welch’s t-test of % responding (ΔF/F·s>10) by trial (2-3 trials per slice; 4 slices from N=4 mice per genotype). (F) Representative UDP calcium response in *P2ry6^-/-^* slice in the absence or presence of bath MRS-2578, a P2Y_6_-selective antagonist (top), and overall quantification (bottom; 4 slices from N=4 mice; unpaired t-test). (G) UDP calcium responses in brain slice at baseline and time points following KA-SE. One-Way ANOVA with Dunnett’s post-hoc comparison to *P2ry6^-/-^* baseline (2 slices/mouse from N=4 mice per group and time point; dot: individual ROI). (H) Microglial process chemotaxis towards a 1mM ATP- or UDP-containing glass pipette. Overall effect of genotype established by Two-Way ANOVA; Slope comparison established by simple linear regression (dotted lines: individual trial mean;). (I) Total distance moved by microglial processes over a 20 min trial. Two-Way ANOVA with Sidak’s post-hoc test (dot: individual process; survey of 28-30 processes from 3 separate slices and animals per group). Bar graphs and shaded line graphs both display the mean ± SEM.

**Figure S6:**
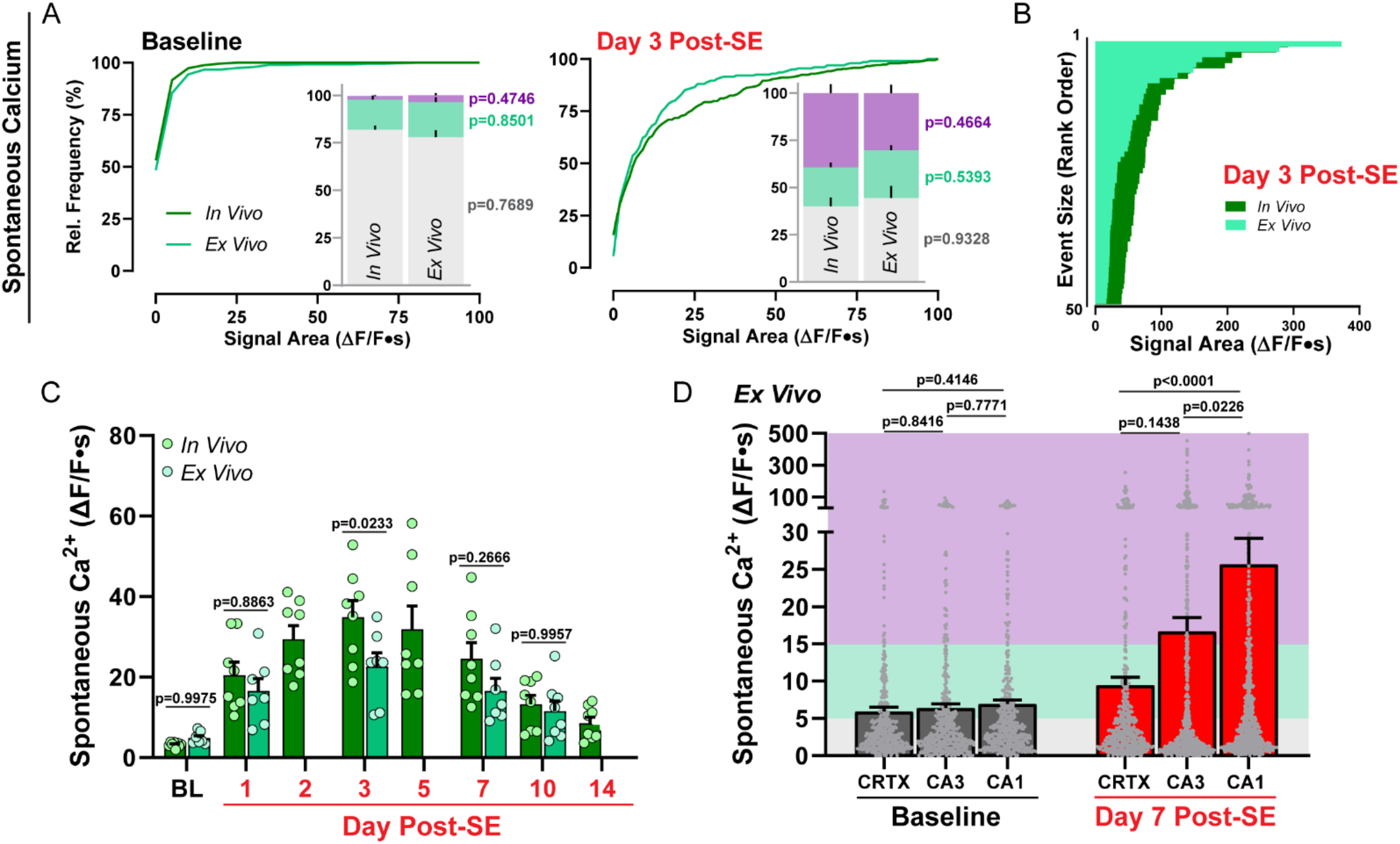
Comparison of spontaneous calcium activity between cortical microglia in slice and *in vivo*, related to Figure 3. (A) Cumulative distribution of spontaneous calcium activity recorded from cortical microglia *in vivo* (chronic cranial window) and *ex vivo* (acute coronal brain slice) at baseline and 3 days after KA-SE. Bar graphs compare the percentage of microglial ROIs which are inactive (gray bar, ΔF/F<5), display low activity (teal bar, ΔF/F=5-15), or are highly active (purple bar, ΔF/F>15). Two-way ANOVA with Sidak’s post-hoc comparison between activity levels. (B) Rank order of the largest calcium events *in vivo* and *ex vivo* in cortex on day 3 after KA-SE. (C) Microglial spontaneous calcium signaling in cortex for *in vivo* and *ex vivo* preparations. Two-way ANOVA with Sidak’s post-hoc test (dot: average spontaneous calcium activity across all ROIs from a chronic window field of view (*in vivo*) or *ex vivo* slice field of view; both are 300 x 300 µm). (D) Comparison of WT microglial spontaneous calcium activity in brain slice by region and time point. One-way ANOVA with Tukey’s post-hoc test by time point (dots represent individual ROIs). Bars represent the mean ± SEM. *In vivo* data come from N=4 WT mice with two regions surveyed longitudinally; *ex vivo* data come from N=4 WT mice per time point with up to two brain slices surveyed per mouse.

**Figure S7:**
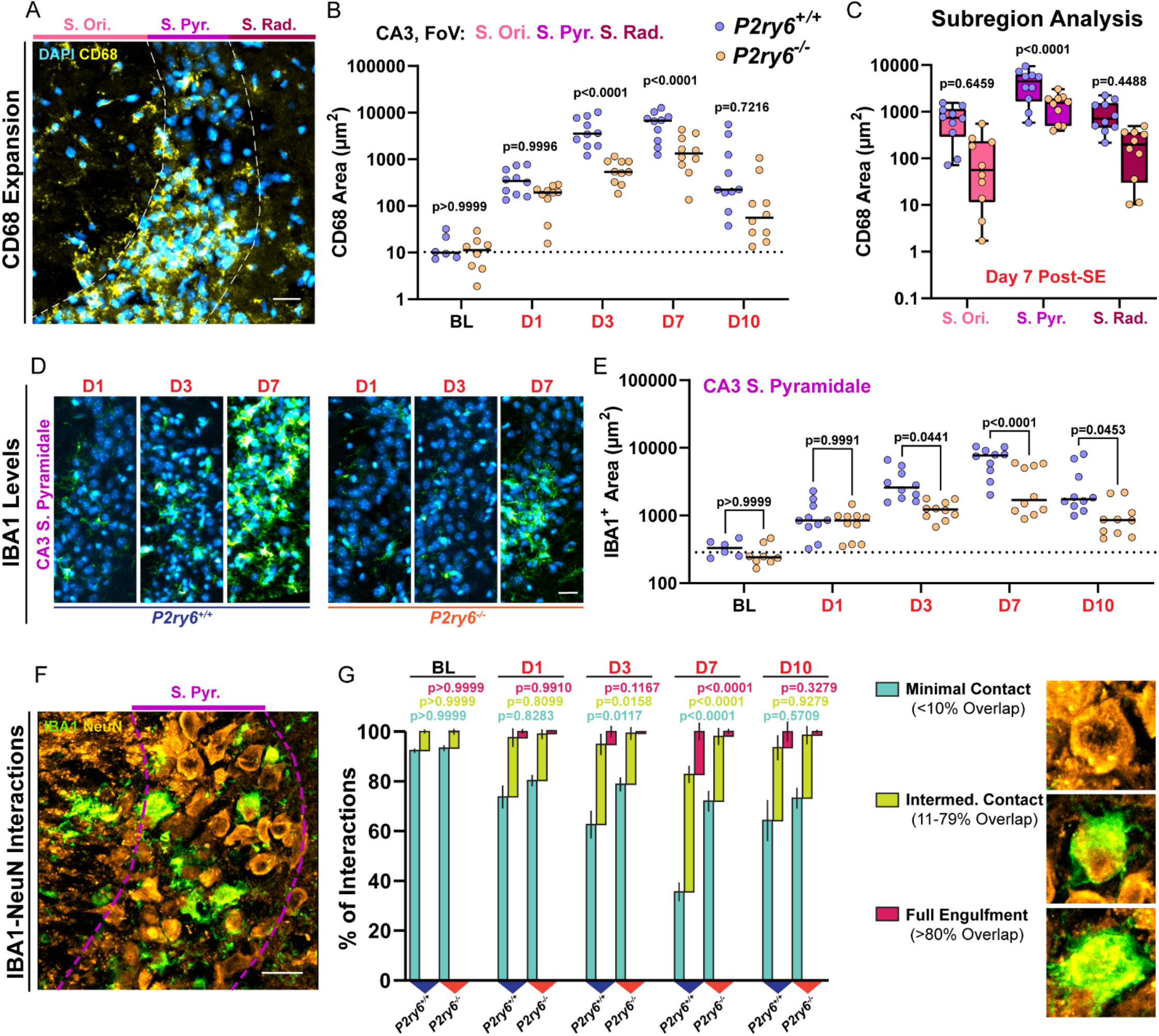
Evolution of phagocytic interactions in CA3, related to Figure 4. (A) Image of CD68 staining in CA3 from a day 7 post-SE WT mouse. Scale bar, 20µm. (B) Quantification of CD68 area over time from the entire CA3 field of view. Two-Way ANOVA with Sidak’s post-hoc test. (C) CD68 staining between genotypes day 7 post-SE by CA3 subregion. One-Way ANOVA with Sidak’s post-hoc test. (D) Representative images of IBA1 (green) in the CA3 pyramidal band during early epileptogenesis. Scale bar, 20µm. (E) Quantification of IBA1 area over time in the CA3 pyramidal band. Two-Way ANOVA with Sidak’s post-hoc test. (F) IBA1 and NeuN cell interactions layer day 7 post-SE in the CA3 pyramidal layer of a WT mouse. Scale bar, 20µm. (G) Percentage of CA3 NeuN neurons having minimal/no contact with IBA1 cells, intermediate contact, or reaching full engulfment criteria across time (examples on the right). Two-Way ANOVA with Sidak’s post-hoc test by interaction type. Bars represent the mean ± SEM. Dots represent one of two hippocampi surveyed per mouse. N=3-5 mice per group.

**Figure S8:**
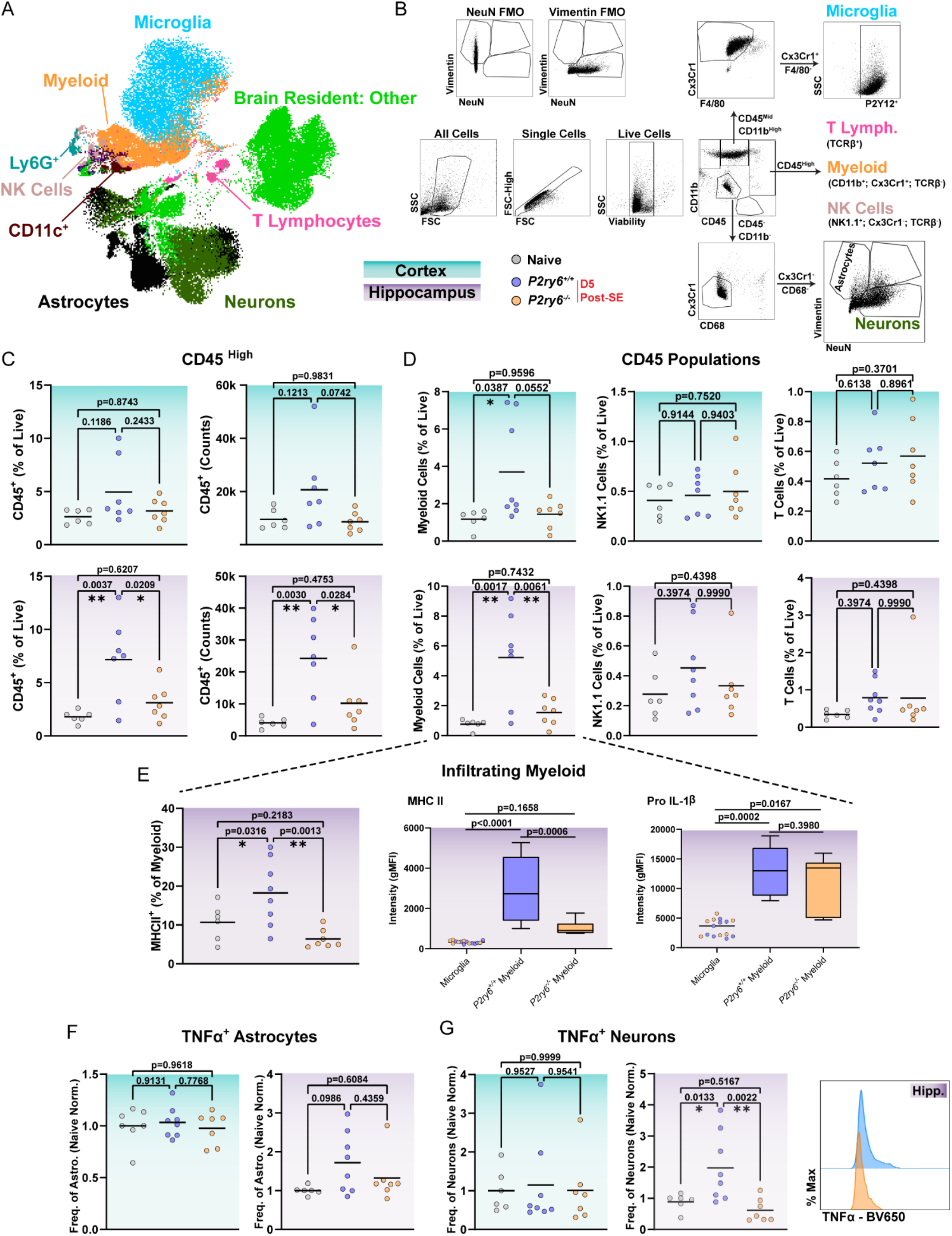
Greater pro-inflammatory myeloid infiltration occurs in *P2ry6^+/+^* hippocampus, related to Figure 6. (A) UMAP of resident and infiltrating cell populations built off FSC, SSC, all surface markers and cytokines (see STAR Methods). (B) Gating strategy to separate microglia, infiltrating immune populations, and other resident brain cells. (C) The number (counts) and frequency (% live) of CD45^+^ immune cell populations between regions and genotypes. (D) Further evaluation of infiltrating CD45^High^ cells based upon markers defining myeloid, NK, or T lymphocyte sub-populations. (E) Differences in MHCII expression frequency and intensity (gMFI) between infiltrating myeloid population in *P2ry6^+/+^* and *P2ry6^-/-^* hippocampus and in reference to resident microglia. Pro IL-1β expression in infiltrating myeloid cells relative to resident microglia. (F) Evaluation of TNFα expression frequency and intensity by vimentin^+^ astrocytes in cortex and hippocampus. (G) Expression of TNFα in NeuN^+^ neurons, including frequency, gMFI intensity, and mode-normalized representative histograms. Data come from N=6 naïve mice (4 WT, 2 KO), N=8 *P2ry6^+/+^*mice and N=7 *P2ry6^-/-^* mice from two independent cohorts. Scatter plots: dot represents one mouse, line at the mean. Box and whisker plots display mean, interquartile range, and min to max. Significance testing utilized a One-Way ANOVA with Tukey’s post-hoc test.

**Figure S9:**
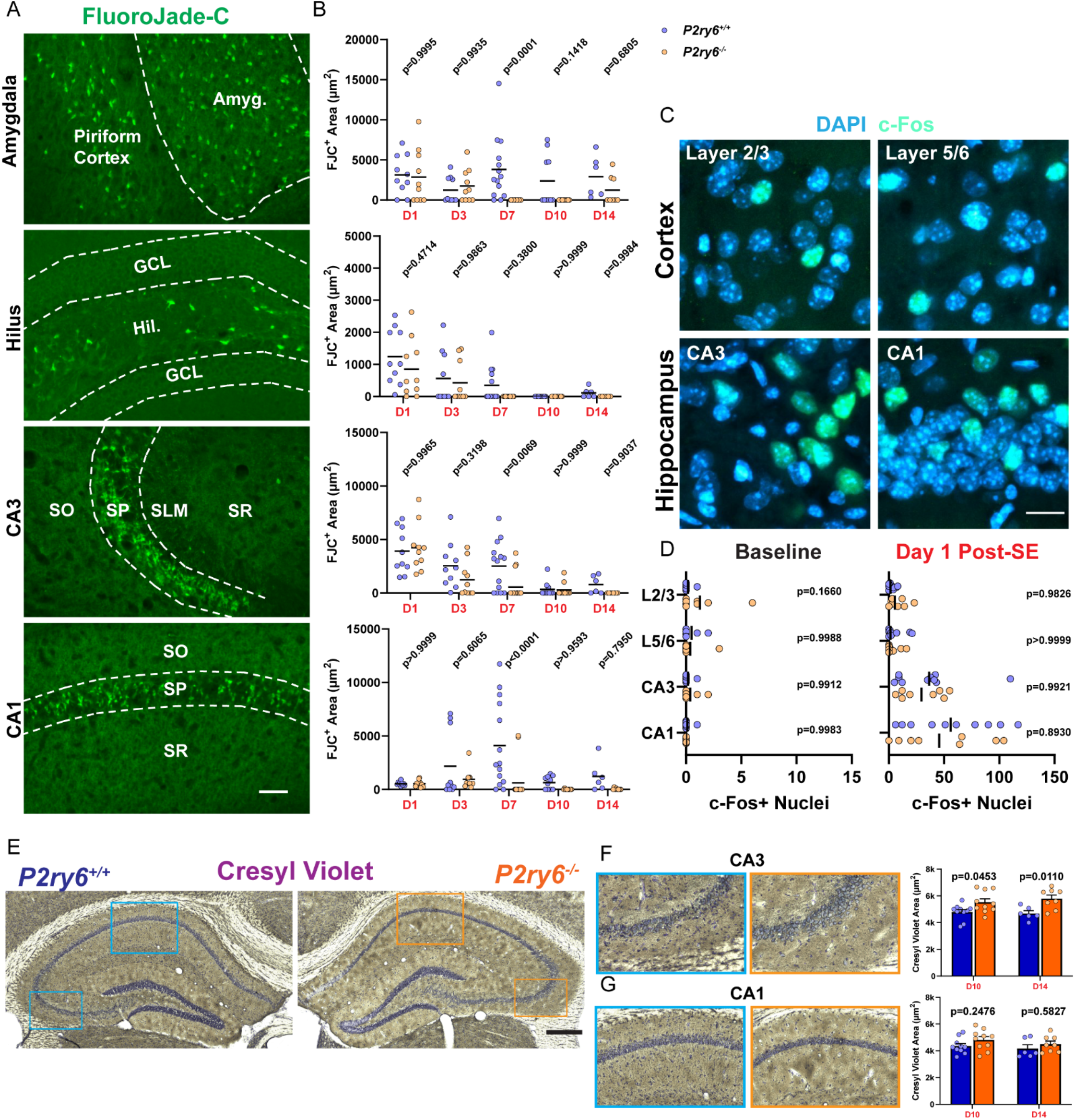
Comparison of neuropathology and seizure damage between genotypes, related to Figure 7. (A) Representative images of FluoroJade-C (FJC) staining one day after KA-SE in the amygdala, hilus, CA3, and CA1 regions. Scale bar, 50 µm. (GCL-Granule Cell Layer). (B) Quantification of FJC-positive cell area by region. Two-Way ANOVA with Sidak’s post-hoc comparison (dot: one region; bilateral survey from N=3-9 mice per group). (C) Examples of DAPI and c-Fos staining one day after KA-SE. Scale bar, 15 µm. (D) Quantification of c-Fos positive nuclei between genotypes at baseline and one-day after KA-SE. Two-Way ANOVA with Sidak’s post-hoc test by region (dot: one region; bilateral survey from N=4-5 mice per group). (E) Representative hippocampi stained with Cresyl Violate acetate 14 days after KA-SE. Scale bar, 500 µm. (F,G) Enlarged images of the CA3 (F) and CA1 (G) regions with corresponding quantification of Cresyl Violet area within the pyramidal band. Two-Way ANOVA with Sidak’s post-hoc test (dot: one region; bilateral survey from N=3-5 mice per group).

## Notes

### Competing Interest Statement

The authors have declared no competing interest.

